# Building pangenome graphs

**DOI:** 10.1101/2023.04.05.535718

**Authors:** Erik Garrison, Andrea Guarracino, Simon Heumos, Flavia Villani, Zhigui Bao, Lorenzo Tattini, Jörg Hagmann, Sebastian Vorbrugg, Santiago Marco-Sola, Christian Kubica, David G. Ashbrook, Kaisa Thorell, Rachel L. Rusholme-Pilcher, Gianni Liti, Emilio Rudbeck, Agnieszka A. Golicz, Sven Nahnsen, Zuyu Yang, Moses Njagi Mwaniki, Franklin L. Nobrega, Yi Wu, Hao Chen, Joep de Ligt, Peter H. Sudmant, Sanwen Huang, Detlef Weigel, Nicole Soranzo, Vincenza Colonna, Robert W. Williams, Pjotr Prins

## Abstract

Pangenome graphs can represent all variation between multiple reference genomes, but current approaches to build them exclude complex sequences or are based upon a single reference. In response, we developed the PanGenome Graph Builder (PGGB), a pipeline for constructing pangenome graphs without bias or exclusion. PGGB uses all-to-all alignments to build a variation graph in which we can identify variation, measure conservation, detect recombination events, and infer phylogenetic relationships.

---

Pangenome graphs compactly represent complete genomes, their homologies (sequence similarities), and all forms of variation between them [1–4]. They allow us to identify variation, measure conservation, detect recombination events, and infer phylogenetic relationships, making them valuable tools for studying sequence evolution and variation in diverse species [5, 6]. However, existing methods for constructing pangenome graphs [7, 8] are biased due to their reference and tree-guided approaches [5, 9], which can lead to incomplete and unstable representations of genetic variation [10]. By adding only sequences that are sufficiently similar to the reference genome, these methods prune regions that are highly variable structurally (section E), such as centromeres and other satellite sequences [7, 8]. Inductive biases result from techniques to mitigate computational complexity [5, 7], or from a goal to structure the resulting graphs so that they are easier to use during read alignment [8]. Although approaches for unbiased whole pangenome graph construction have been proposed and are capable of scaling to whole genome analyses [10–12], these often lead to complex and difficult-to-use graph models or introduce new complexity for downstream analyses due to the effects of the specific k-mer length and other parameters [12]. Projects which build resources on tree and reference-based alignment methods [5, 13] would benefit from the availability of unbiased, reference-free whole genome alignments to control the quality of their results.

To overcome these limitations, we propose the PanGenome Graph Builder (PGGB), a reference-free pipeline to construct unbiased pangenome *variation graphs* [10]. Its output presents a base-level representation of the pangenome, including variants of all scales from single nucleotide polymorphisms (SNPs) to structural variants (SVs). The constructed graph is unbiased, i.e., all genomes are treated equivalently, regardless of input order or phylogenetic dependencies, and lossless: any input genome is completely retained in the graph and may be used as a frame of reference in downstream analysis. PGGB makes no assumptions about phylogenetic relationships, orthology groups, or evolutionary histories, allowing data to speak for itself without risks of implicit biases that may affect inferences made from the graph. PGGB is implemented as a modular shell script, integrating independent components via standard text-based file formats, which provides a template for future pangenome construction methods. The method, developed and applied over several years within the Human Pangenome Reference Consortium (HPRC) [18, 19], has proven to be accurate and scalable to hundreds of genomes, as confirmed by the broader bioinformatics community [20–22]. Here, we describe the specific innovations in the three main phases of the algorithm: alignment, graph creation, and graph normalization. We then use cross-validation with MUMMER4 [23] to demonstrate the accuracy of our approach across a wide range of species that differ greatly in genome variation and scale (Table 1). In cases with suitable data, we compare genome variation captured by the graphs to variants identified by conventional reference-based methods (Section B.5). Finally, we study the openness of constructed pangenomes, a biological parameter which describes the degree to which new genomes are expected to contribute new sequence (F).

**Table 1:** Performance of PGGB with pangenomes across species. For each pangenome, we report its size, the compression ratio (pangenome sequence length divided by graph size), the PGGB runtime, the maximum memory usage of PGGB, the average number of SNVs (across all haplotypes except the one used as reference) identified with MUMMER4 that we used to evaluate SNVs identified with PGGB, and the average F-score (across all haplotypes except the one used as reference) computed using MUMMER4’s SNVs as ground truth. The name of each pangenome indicates the species and the number of haplotypes. All runs were performed on machines equipped with AMD EPYC 7402P 24-Core, 378 GB of RAM, and a 1 TB Solid-State Drive. All PGGB runs were executed with 48 threads. *Erdős–Rényi random sparsification activated.

| Pangenome | Size (bp) | Compr. | Time (h) | Mem. (GB) | N(SNVs) | F-score |
| --- | --- | --- | --- | --- | --- | --- |
| athaliana82 | 11052471644 | 36.28 | 204.06 | 130.59 | 572924 | 0.920053 |
| ecoli50 | 255615162 | 11.50 | 0.93 | 19.93 | 56296 | 0.977863 |
| ecoli500* | 2562798947 | 23.75 | 41.39 | 210.87 | 60243 | 0.967453 |
| hsapiens90.chr6 | 15508376475 | 86.88 | 18.46 | 137.72 | 147580 | 0.975822 |
| mouse36.chr19 | 2150650973 | 11.68 | 3.80 | 28.66 | 166734 | 0.940177 |
| primate16.chr6 | 2661626374 | 8.12 | 5.60 | 37.05 | 2309269 | 0.947453 |
| scerevisiae142 | 1702093905 | 57.12 | 20.46 | 119.68 | 61439 | 0.971758 |
| scerevisiae142* | 1702093905 | 45.17 | 10.11 | 80.25 | 61232 | 0.972030 |
| soy37 | 36673233537 | 21.51 | 99.73 | 37.54 | 1302099 | 0.942402 |
| tomato23 | 18691283883 | 19.73 | 22.34 | 42.06 | 861654 | 0.967951 |

PGGB begins with sequence alignment (Figure 1A). The method supports any type of genomic sequence data in FASTA/Q file format (sequencing reads, genes, genomes, or a combination of these). To avoid reference and order bias, we use an all-to-all alignment of the input sequences. This approach aligns sequences directly to each other, enabling each sequence in the pangenome to serve as a potential reference to describe variation. Problematically, this requires all-to-all comparisons that scale quadratically with the number of included genomes [10]. For this reason, PGGB uses WFMASH [24], which reduces costs by mapping and aligning in the space of sequence segments rather than single base pairs. WFMASH first applies an extension of MASHMAP [25] to obtain homology mappings, by default using seeds of 5 kbp to find similarities of 25 kbp or more at 90% average nucleotide identity. WFMASH then uses a generalization of the bidirectional Wavefront Algorithm (BiWFA) [26, 27] that aligns the sequences by comparing segments of 256 bp rather than single characters. This algorithm, BiWF*λ*—so named because it replaces character match with a callback function *λ* that matches segments—obtains a final base level alignment by splicing together “incepted” alignments over the 256 bp segment pairs that lie in the optimal alignment path. Our use of WFMASH ensures that the alignments that structure the graph feature long-range collinearity that is insensitive to repetitive, shorter similiarites found between transposons and satellite sequences. Although WFMASH alignments have an ideal structure for its operation, PGGB can build the graph using any set of user-defined alignments in PAF format.

**Fig. 1:**
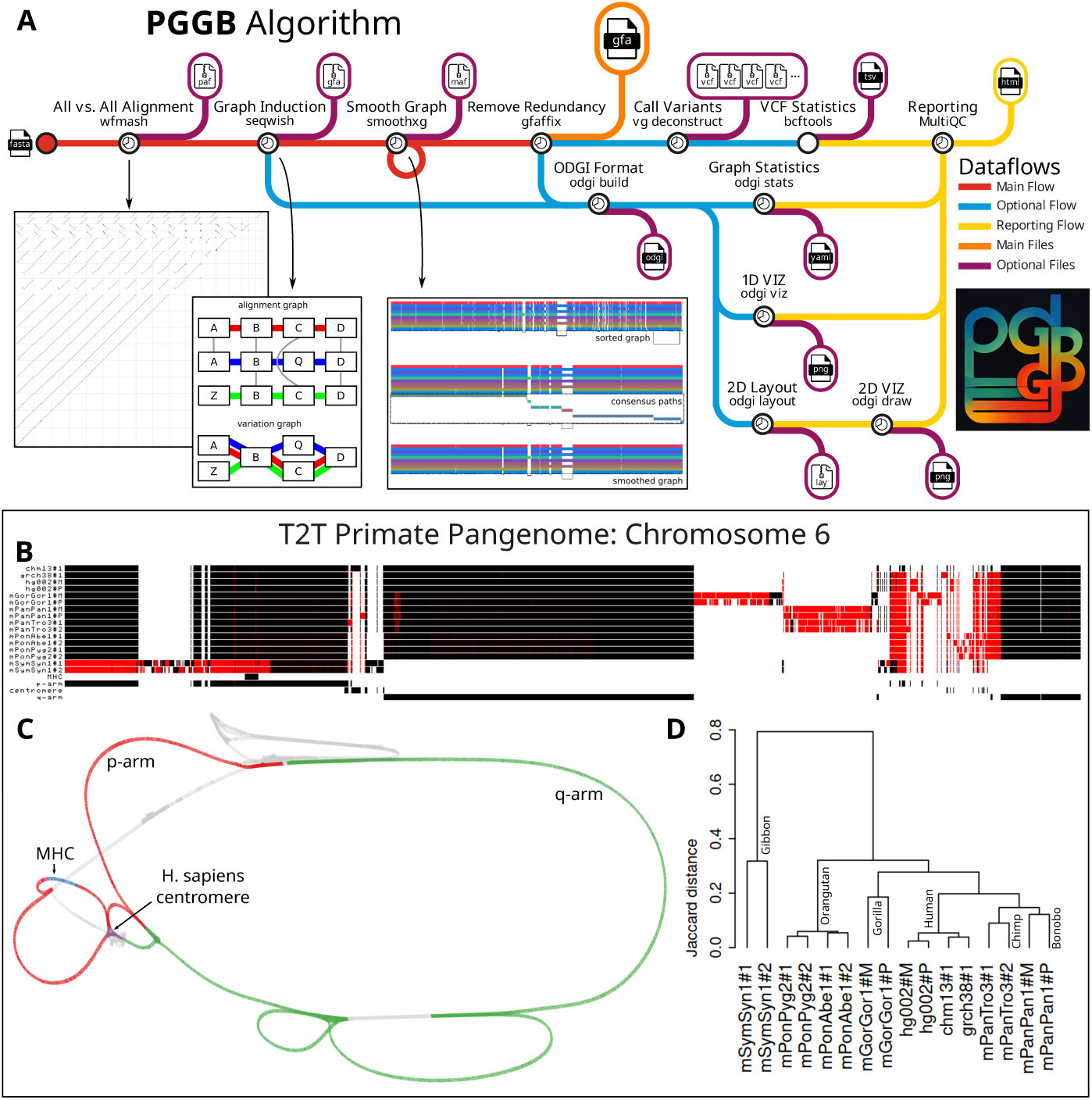
PGGB and applications. (A) PGGB’s algorithms/data flows. Primary flow (red) proceeds from FASTA to alignment, graph induction, smoothing, to normalization with GFAFFIX [14], ending with the final variation graph (orange). Optional outputs (blue): statistics, variant calls, and 1D/2D graph visualizations. (B) 1D pangenome graph visualization using 16 haplotype-resolved primate assemblies homologous to human chromosome 6. T2T-CHM13 annotations (Major Histocompatibility Complex, p-arm, centromere, q-arm) are shown. Black indicates that the path traverses the graph in the forward orientation, while red indicates that it traverses in the reverse. The p-arm region with the MHC is inverted in gibbon. Centromeric regions appear largely dissimilar among many species, except between chimpanzee and bonobo, and between the orangutans. (C) 2D visualization, rendered with the same human chromosomal annotations in GFAESTUS [15], shows possible circularization of the graph due to structural variation or homogenization between subtelomeric regions. (D) Using ODGI [6], we extract a pairwise distance matrix based on in-graph Jaccard metrics over shared base pairs. This distance matrix yields a phylogenetic tree that matches previous results based on SNPs [16]. We posit that the greater phylogenetic distances reflect the inclusion of the centromeres—which feature low rates of recombination and tend to diversify rapidly by near-clonal evolution [17]—in our distance computation.

The second step—pangenome graph induction—converts a collection of genomes and pairwise alignments into an equivalent variation graph. We achieve this with SEQWISH [10], a tool specifically designed to scale graph induction to whole pangenomes in low memory. At a high level, SEQWISH merges all DNA base-pairs that are matched together in the alignments into a single node in the output graph. This process also compresses transitively-matched base-pairs. For example, if *A*, *B*, and *C* are characters in input sequences and → represents a character match, *A → B → C* would result in a single node in the output graph that also implies the transitive match *A → C*. Each input genome is then fully embedded in the graph as a path, recording graph edges where nodes occur successively in the path.

The SEQWISH graph recovers transitive homology relationships that may not be present in the initial alignment set (Figure 1A). This property allows us to apply random sparsification to reduce the complexity of very large alignment problems. We first compute the all-to-all homology mapping with WFMASH, which is quadratic but highly efficient due to its use of sketched representations of the mapped sequences. To greatly reduce runtime for large numbers of input genomes, we only compute the pairwise alignments for a random subset of the mappings. Our goal is to retain enough mappings that we expect the alignment graph to be fully connected locally at each homologous locus in the pangenome. To determine a safe threshold for sparsification, PGGB uses a heuristic based on the Erdős–Rényi random graph model. For a random graph with *N* nodes, this model predicts that as *N → ∞* the graph is almost certainly fully connected so long as pairs of nodes are connected with probability 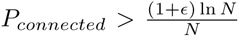 (where *ɛ* is a small constant) [28]. We thus set a sparsification parameter that uses a hash of each mapping record to drop mappings with a probability *P_sparse_* ≫ *P_connected_* to ensure that while some edges are removed, connected component encompassing critical homologous relationships is preserved. This lets us reconstruct all transitive relationships in the variation graph without needing to directly compute all pairwise alignments, avoiding the expected *O*(*N*^2^) costs implied when *P_sparse_* = 1. This dramatically reduces the runtime of alignment and graph induction with negligible effect on accuracy (Table 1), e.g. 10× increase in the number of genomes requires only 44× increase in runtime—rather than 100 ×.

Graph building completes with SMOOTHXG (Figure 1A), an iterative post-processing step specifically designed for PGGB that locally compresses and simplifies the pangenome variation graph. Although the SEQWISH graph presents a complete, lossless model of the input genomes and their homologies, in our experience it often presents complex local motifs that can cause problems for diverse types of downstream analysis (Section B.2). A key issue is that pairwise alignments derived across input sequences are not mutually normalized, leading to different representations of small variants like indels in low-complexity sequences, which in turn generate complex looping motifs that are difficult to process. We mitigate this issue by removing short matches from SEQWISH’s input alignments. This reduces complexity, but also creates a graph that can be locally “under-aligned” and does not represent all local pairwise relationships. To resolve this, we apply a local realignment kernel, partial order alignment (POA) [29–31], across all parts of the graph. By default, we do so at a scale of around 1kbp, which is smaller than most nonlinear patterns of structural variation found in genomes [18, 32]. This allows the PGGB graph to represent complex structurally-variable loci as simple loops through a single copy of duplicated sequences [18]. The kernel is applied to regions that are extracted from a 1-dimensional graph embedding [6, 33]. This embedding orders nodes in the graph so that their distance in the order best-approximates their distance in the genomic paths of the graph. SMOOTHXG first learns this embedding, then obtains partially overlapping segments of the graph (blocks) to which it then applies POA. The realigned blocks are “laced” back into a complete variation graph. We iterate the entire SMOOTHXG step multiple times (3 by default) to limit edge effects that can occur near block boundaries, progressively refining the learned graph embedding. As a final normalization step, we apply GFAFFIX to compress redundant nodes [14] and use ODGI to make a final sort for the modified graph [6].

PGGB provides outputs that support immediate interpretation, quality control, and downstream applications. Using ODGI, it produces basic graph statistics, such as size, node count, and base content. ODGI creates 1D and 2D visualizations that provide intuition about the structure of the entire graph, with the 1D view showing the relative alignment of paths into the graph structure, and the 2D view showing high-level features of the graph topology. Both can be applied at the scale of multi-gigabasepair graphs. Optionally, PGGB provides graph statistics and diagnostic plots in a MultiQC report [34]. We also provide an option to call variants [1, 35] from the graph to produce a phased description of embedded haplotypes in variant call format (VCF). Variants called directly from the graph can include large nested genetic sites, leading to incompatibility with many applications. To address this, PGGB decomposes complex nested variation into a minimal reference-relative representation using BiWFA [36]. This allows PGGB to provide input to analyses based on small variants, leading to compatibility with numerous downstream applications based on genomic variation. PGGB is thus a multi-sample variant caller for whole-genome assemblies.

PGGB has been applied and validated at large scale in projects in the HPRC [18], where it additionally has provided the first sequence-based evidence for systematic recombination between heterologous acrocentric human chromosomes [19]. Here, to demonstrate PGGB’s broad utility, we present results from its application to a variety of diverse pangenome and comparative genomic contexts (Table 1, Figure 1, Supplementary Figures F13, B4, B5, B6, D7, F11, F12, F13). We provide information on runtime and resource requirements, showing that even for hundreds of small genomes, PGGB can provide a variation graph within hours. Larger genomes in general require partitioning to maximize parallelism and ensure that total compute requirements fit in standard commodity servers (Section B.3). Due to lack of ground truth, quality evaluation on real data can be difficult. For validation, we compare PGGB’s output with SNPs detected by MUMMER4 [23], a current standard approach for pairwise whole genome alignment. These show cross validation F-scores *>*92% across all tested contexts, indicating that the method performs equivalently to existing standards. The difference between PGGB and MUMMER4 is driven by different alignment representations in regions with very close SNVs (Section G). However, while MUMMER4 provides only pairwise comparisons with a target reference, PGGB yields a full all-to-all comparison between genomes that leads to completely new bioinformatic analysis modalities.

Many downstream applications that are typically based on polarization of variants (e.g. SNPs) relative to a single reference genome may be directly implemented in the space of variation graphs built with PGGB and similar methods. This follows from two basic concepts: in the variation graph, nodes are alleles, while genomes can be simply understood as vectors of allele counts. Methods based in this vector space allow us to simultaneously consider *all* classes of variation in downstream analyses, without reference bias. As proof of principle, we put forward a phylogenetic tree constructed directly from distances measured within a pangenome variation graph of 16 complete assemblies of chromosome 6 from the great ape family (Figure 1D), which matches established phylogenies of the *Hominoidea* clade based on manually curated sets of SNPs that exclude structurally variable regions [16]. We have also successfully applied this method in yeast, where a comparison of phylogenetic trees built from MUMMER’s SNVs against the reference and PGGB graph’s nodes showed that the latter correctly infers the phylogeny in a number of cases where both haplotypes from the same genome are brought together (Section D). We also noticed that the same approach when applied to the primate graph made with Minigraph-Cactus led to a different phylogenetic tree (E9).

PGGB is a new, modular, and conceptually straightforward approach to understanding sequence relationships between many complete genomes in both pangenomic and comparative genomics settings. Our approach provides a general framework for genome graph building techniques which we expect researchers will upgrade and extend in the future. A cluster-scalable Nextflow implementation is already in progress [37]. By making it easy to build variation graphs, PGGB opens the door to diverse downstream population and evolutionary genetic methods that can consider all classes of sequence variation simultaneously. This will allow us to develop a comprehensive understanding of the links between sequence variation, phenotype, and evolution in an era where the complete assembly of genomes becomes routine.

**Online content.** PGGB is available at https://github.com/pangenome/pggb. Code used for experiments can be accessed at https://github.com/pangenome/pggb-paper. Pangenomes are available at https://doi.org/10.5281/zenodo.7937947.

## Acknowledgments

The authors thank members of the HPRC Pangenome Working Group for their insightful discussion and feedback, and members of the HPRC production teams for their development of resources used in our exposition.

## Author contributions

*Project conception*: Erik Garrison

*Project guidance*: Erik Garrison, Sven Nahnsen, Nicole Soranzo, Vincenza Colonna, Robert W. Williams, Pjotr Prins

*Software development* : Erik Garrison, Andrea Guarracino, Simon Heumos,

Santiago Marco-Sola, Mwaniki N. Moses

*Paper editing*: Erik Garrison, Andrea Guarracino, Simon Heumos, Vincenza Colonna, Robert W. Williams, Pjotr Prins

*Experimental design*: Erik Garrison

*Quality evaluation*: Erik Garrison, Andrea Guarracino, Lorenzo Tattini

*Testing* : Erik Garrison, Andrea Guarracino, Simon Heumos, Flavia Villani, Zhigui Bao, Lorenzo Tattini, Jörg Hagmann, Sebastian Vorbrugg, Christian Kubica, Kaisa Thorell, Rachel L. Rusholme-Pilcher, Agnieszka A. Golicz, Sven Nahnsen, Zuyu Yang, Mwaniki N. Moses, Franklin L. Nobrega, Hao Chen, Joep de Ligt, Peter H. Sudmant

*Experimental execution*: Andrea Guarracino

*Documentation*: Andrea Guarracino, Simon Heumos

*Mus musculus and Rattus Norvegicus*: Flavia Villani, David G. Ashbrook, Hao Chen, Vincenza Colonna

*Tomato pangenome*: Zhigui Bao, Sanwen Huang

*S. cerevisiae* and *S. paradoxus* : Lorenzo Tattini, Gianni Liti

*Soy G.max* : Jörg Hagmann

*A. thaliana*: Sebastian Vorbrugg, Christian Kubica, Zhigui Bao, Detlef Weigel

*Parameter settings*: Sebastian Vorbrugg

*Algorithm development* : Santiago Marco-Sola

*Helicobacter pylori* : Kaisa Thorell, Emilio Rudbeck

*V. fava*: Agnieszka A. Golicz

*Neisseria mingitidis*: Zuyu Yang, Joep de Ligt

*SARS-CoV-2 dataset* : Mwaniki N. Moses

*E. coli and Coliphages*: Franklin L. Nobrega, Yi Wu

*Primate pangenome*: Peter H. Sudmant

*High Performance Computing management* : Pjotr Prins

## Appendix A Data

Lists of all accessions for all pangenomes are reported in the Supplementary File 1.

### A.1 A. thaliana

We downloaded the assemblies from Genbank [38], considering those that resolved the genome with 5 contigs, one for each chromosome, and removing the contigs representing mitochondria and plasmids. Furthermore, we also included GCA 028009825.1, obtaining the final set of 82 assemblies.

### A.2 E. coli

We downloaded the assemblies from Genbank [38], considering those that completely resolved the genome. From these, we randomly selected 500 and 50 assemblies.

### A.3 H. sapiens

We downloaded the assemblies from [18], considering contigs belonging to chromosome 6, obtaining the final set of 90 haplotypes (from 47 individuals plus two reference genomes, that is CHM13 and GRCh38).

### A.4 M. musculus

We downloaded the assemblies from Genbank [38], considering those that resolved the genome at chromosome level, obtaining the final set of 36 assemblies.

### A.5 Primates

We downloaded the assemblies for *Gorilla gorilla*, *Pan paniscus*, *Pan troglodytes*, *Pongo abelii*, *Pongo pygmaeus*, *Symphalangus syndactylus* released by the Telomere-to-Telomere consortium. To this set, we also included 3 human genomes, that is CHM13 and GRCh38, which are haploid, and both MATERNAL and PATERNAL haplotypes of the HG002 v1.0.1 assembly, totaling a final set of 16 haplotypes.

### A.6 S. cerevisiae

We downloaded 142 assemblies from [39].

### A.7 Soybean

We downloaded 38 assemblies from [40] and [41], covering the following species:

*Glycine soja*, that is wild soybean, and *Glycine max*, that is cultivated soybean.

### A.8 Tomato

We downloaded 23 assemblies from [42], covering the following species: *Solanum lycoperscium*, *Solanum pimpinellifolium*, and *Solanum lycopersicum var. cerasiforme*.

## Appendix B Methods

Here we provide details about components which are not described in other publications. Our primary focus is on SMOOTHXG, the algorithm that PGGB uses to locally normalize and simplify the graph produced by previous phases (alignment with WFMASH and graph induction with SEQWISH). Through a series of passes over the pangenome, SMOOTHXG reshapes the graph to reduce local complexity and underalignment. This resolves key problems encountered in earlier attempts to implement all-vs-all alignment based graph construction [10, 43], which typically resulted in very complex, looping, graph motifs at small scales, and graph redundancy or loss of alignment sensitivity caused by match filtering. We additionally describe the evaluation method we use in our cross-validation experiments where PGGB graphs are compared with SNPs determined by MUMMER4.

### B.1 SMOOTHXG

SMOOTHXG requires a GFA pangenome graph as input, for example, the output from SEQWISH. The raw SEQWISH graph is globally unsorted and might be locally unaligned. SMOOTHXG sorts and normalizes the graph preserving nonlinear complex global structural variation. Detailed steps are described subsequently.

#### Preprocessing

A Path-Guided Stochastic Gradient Descent (PG-SGD) algorithm optimizes the one-dimensional (1D) node order of the graph to best match the nucleotide positions in the embedded paths. A grooming step ensures that for each node, the node orientation follows the consensus node orientation of all path steps visiting the node. A 1D topological sorting of the graph completes the overall sorting steps. Finally, the graph is chopped so that each node contains a relatively small number of base pairs (SMOOTHXG default: 100 bp), preserving node topology and order. Node chopping prepares the graph for more exact cut points during the block creation process described in the next section. Since blocks are constrained by a maximum length, without cutting nodes, having long nodes could lead to the formation of large blocks that do not properly represent the local variation of all genomes in the graph. To draw an analogy, consider the task of evenly distributing smaller pieces of an object into various containers. Smaller parts are inherently easier to allocate properly. Similarly, shorter pieces of sequences (the pangenome graph nodes) are more conveniently distributed into blocks, ensuring a more uniform representation of the genomes in the graph. However, nodes that are too small increase the requirements for keeping the graph in memory and working with it. Chopping nodes to have a maximum length of 100 bp is a trade-off between creating uniform blocks and the hardware requirements for working with the chopped graph.

#### Create blocks

The smoothable blocks are discovered by iterating over all nodes following the previously calculated order. A node is added to a block if its addition does not exceed the: 1. total path length limit of a block, 2. the maximum edge jump limit of a block, or 3. the maximum block length. Blocks are broken at likely Variable Number Tandem Repeat (VNTR) boundaries. The VNTR detection is based on a statistical analysis using the autocorrelation. Autocorrelation (aka serial correlation) measures the similarity of a sequence with a lagged version of itself. When applied to nucleotide sequences, this helps to identify repetitive regions. This approach is probabilistic and relies on statistical thresholds to identify potential VNTRs. Here is a breakdown of the key steps:

1. **Autocorrelation calculation:** We calculate the autocorrelation of the nucleotide sequence for different lag values (i.e., lengths of the repeat unit).
2. **Statistical analysis:** We calculate the mean and standard deviation of the computed autocorrelation values. Then, we transform these into Z-scores, which are measures of how far and in what direction individual observations are from the mean, expressed in units of standard deviation.
3. **Identification of likely VNTRs:** We identify putative VNTRs by looking for peaks in the Z-scores. A peak in the Z-scores indicates a lag length where the autocorrelation is significantly higher than average, indicating a strong repetitive pattern, then suggesting a repetitive region.

Furthermore, blocks are broken to ensure that the path ranges within each block do not exceed the maximum sequence input size for the POA step described in the next section. By default, we do so at a scale of around 1kbp, which is smaller than most nonlinear patterns of structural variation found in genomes.

#### Smooth each block

For each block, padding extends each block to the left and right. This improves the local alignment at the boundaries of each block. The k-mer plus min-hash approach ensures that only unique sequences are passed to the POA step, which can significantly reduce runtime. POA is applied to each block, applying by default the following scoring model: 1 for matches, 19 for mismatches, 39 as gap opening penalty, 3 as gap extension penalty, 81 as gap opening penalty of the second affine function, 1 as gap extension penalty of the second affine function. Such scores are tuned for low divergence sequence alignments, as blocks are built to have locally similar sequences. Optionally, this step generates a multiple sequence alignment in MAF format for each block. After the sequence alignment, the padding is removed (block trimming), and then the block is saved for later integration into a full graph.

#### Lace blocks into the smoothed graph

The smoothed blocks are laced together to the final pangenome graph following their initial input order. As a final step, the graph is unchopped to restore the maximum possible node lengths in the graph.

**Fig. B1:**
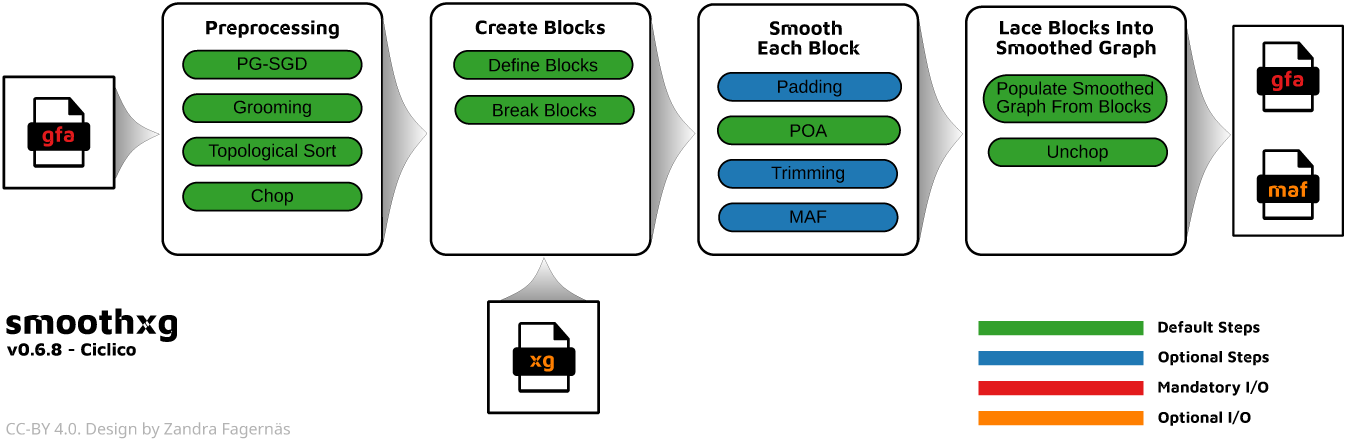
Overview of the algorithmic steps in SMOOTHXG.

### B.2 Graph normalization

SMOOTHXG is specifically designed to locally compress and simplify the raw SEQWISH variation graph which often presents complex local motifs that increase graph complexity. To explicitly demonstrate its efficacy, we built pangenome graphs representing 90 human chromosome 6 with and without applying SMOOTHXG. The graph obtained without SMOOTHXG, despite having fewer nodes than the graph obtained by applying SMOOTHXG (209420591 versus 212423296), has regions where the depth and degree (number of edges) of the nodes are significantly greater (Figure B2). These regions, typically centromeres and other satellite sequences, cause problems for different types of downstream analyses, hindering the performance of all possible operations applied to the graph.

**Fig. B2:**
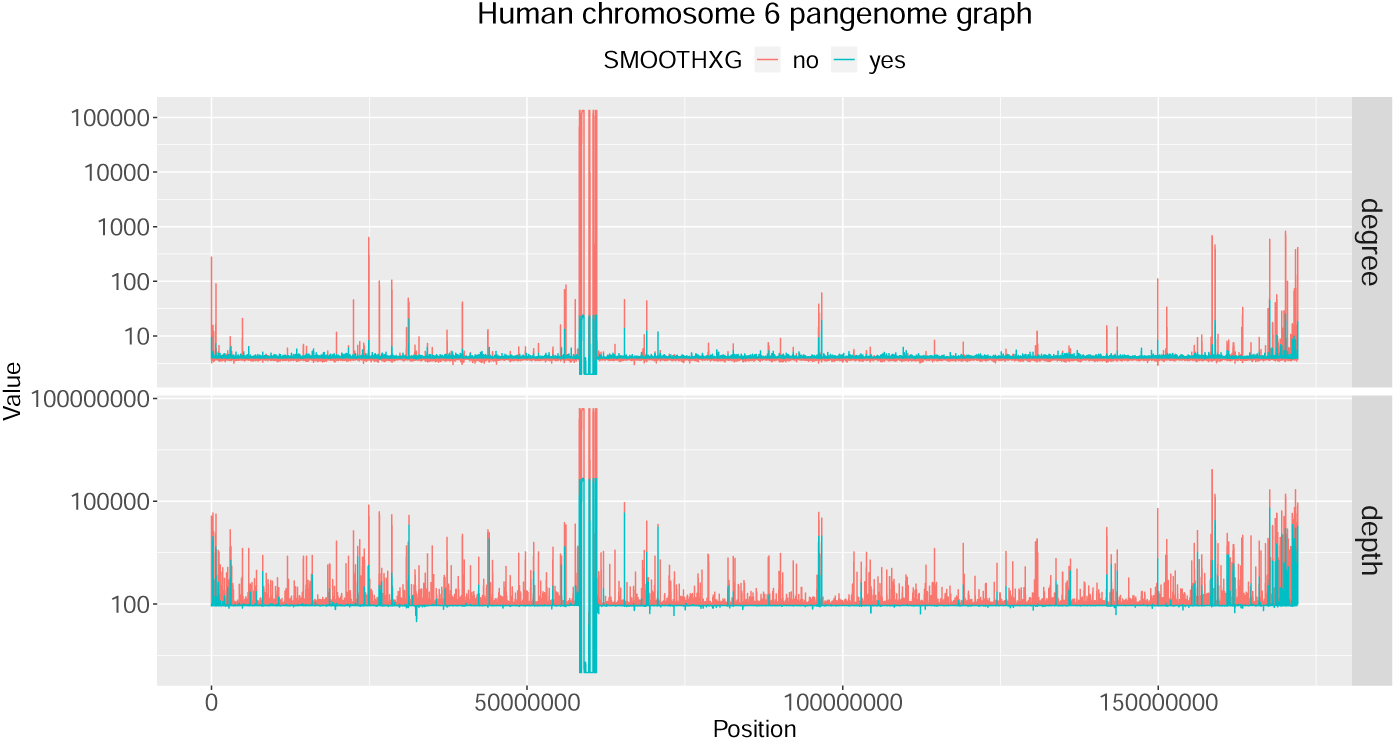
SMOOTHXG locally compresses and simplifies the raw SEQWISH’s graph.

Importantly, SMOOTHXG applies a series of graph sorting algorithms that sort the graph node in a way that better reflects the genome variation embedded in the graph. As an example, we report a representation of the complement component 4 (C4) subgraph extracted from the chromosome 6 pangenome graph (Figure B3). We only show 8 haplotypes (GRCh38, CHM13, and 3 diploid individuals) for simplicity. Figure B3a represents the C4 graph extracted from the pangenome graph built without SMOOTHXG. Node order is not optimal, so the underlying genome variation is still not clearly visible. By applying to such a graph several graph sorting algorithms (implemented in ODGI and applied in SMOOTHXG), the visualization in Figure B3b improves, but we still have a complex graph topology which is difficult to sort cleanly, due to local motifs that are often present in the raw SEQWISH’s graph. Figure B3c represents the C4 graph extracted from the chromosome 6 pangenome graph built with SMOOTHXG. Both node order and graph topology properly represent genome variation present in such a region.

**Fig. B3:**
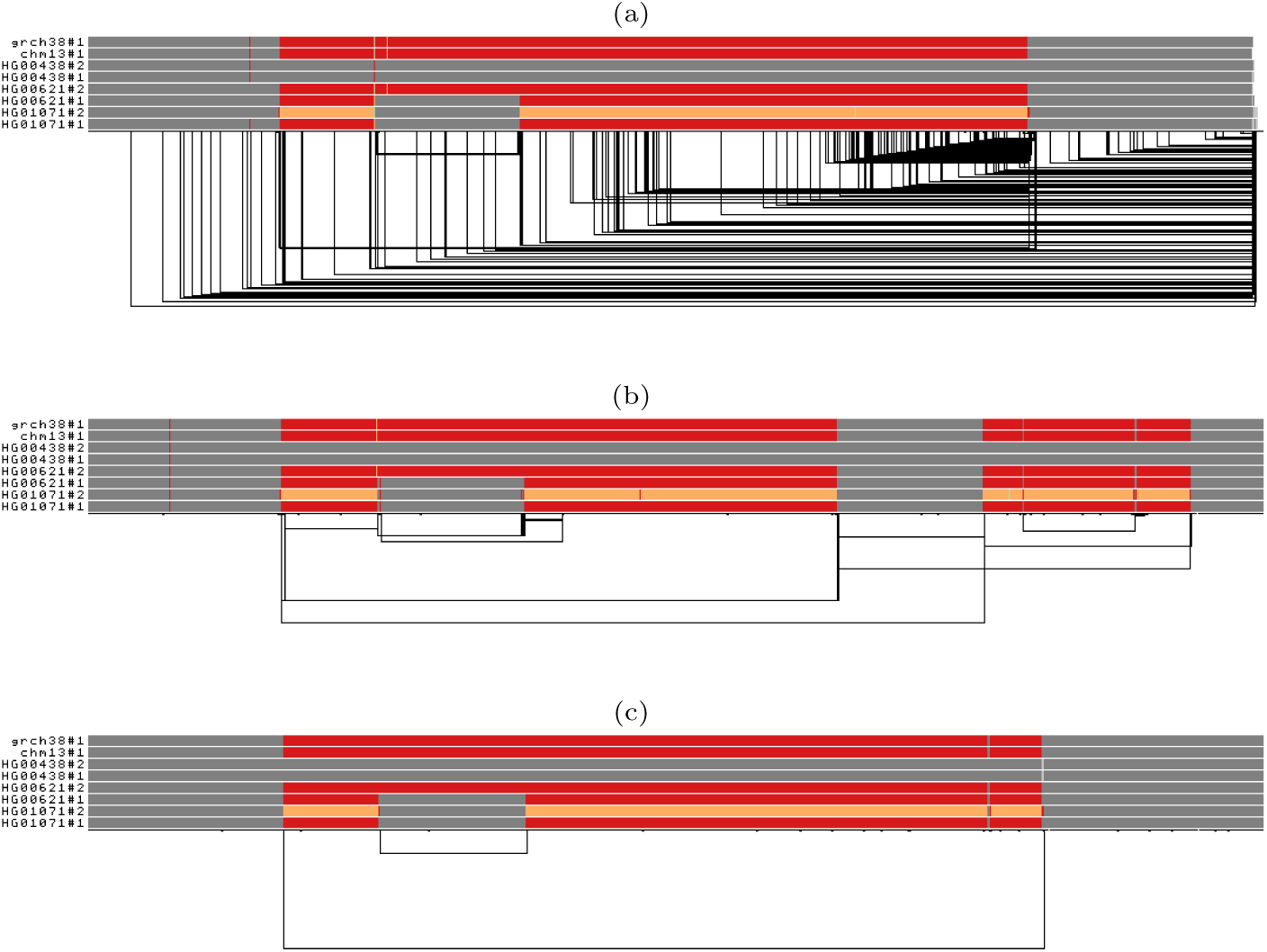
Graph normalization effect on complement component 4 (C4) pangenome graph complexity. Each bar represents a haplotype and black lines on the bottom represent the graph topology. Paths are colored by using the Spectra color palette with four levels of node depths: white indicates no depth, while gray, red, and yellow indicate depths 1, 2, and greater than or equal to 3, respectively. a) C4 subgraph extracted from the chromosome 6 graph built without SMOOTHXG. b) C4 subgraph extracted from the chromosome 6 graph built without SMOOTHXG and sorted. c). C4 subgraph extracted from the chromosome 6 graph built with SMOOTHXG. The two references present two different allele copies of the C4 genes (red = 2X coverage), both of them including the HERV sequence. The entirely gray paths have one copy of these genes (grey = 1X coverage). HG01071#2 presents three copies of the locus (orange = 3X coverage), of which one contains the HERV sequence (gray in the middle of the orange).

### B.3 Sequence partitioning

Pangenome graphs can represent the mutual alignments within a collection of sequences. Due to dispersed sequence similarities caused by transposons, copy number variants, and segmental duplications, a clean partitioning of sequences into isolated components is typically not possible. However, breaking the problem of building the graph into smaller parts is important for reducing the difficulty of computing the pangenome graph. This leads us to need to partition the construction problem. To do so, before applying PGGB, we can perform community detection with the input sequences to uncover the underlying structure of their mutual relationships and gather group of sequences that share common properties. For example, the identified communities (or groups of sequences) may correspond to the distinct chromosomes present within the input genomes. By incorporating reference genomes into the set of input sequences, we can leverage the reference annotations to effectively partition sequences by chromosome. Once these groups of sequences have been identified, we can then partition the sequences by community and apply PGGB on each group separately.

To simplify the sequence partitioning process for users, we provide such a workflow as a dedicated shell script called “partition-before-pggb”. Here is a summary of its key steps:

1. **Homology detection.** We perform pairwise mapping of the input sequences with WFMASH [24] to identify homologous regions, specifically focusing on their location, size and estimated identity. We do not require base-level alignment; by omitting this step, we accelerate the homology detection.
2. **Mapping graph construction.** We use the homology map to build a mapping graph. In contrast to a variation graph, each node in a mapping graph represents an input sequence, with edges representing mappings between these sequences. The edges are weighted, with each weighting factor given by the product of the length of the mapping and its estimated sequence identity.
3. **Community detection.** We apply the Leiden algorithm [44] to detect the underlying communities present in the mapping graph. In the context of community detection, a community is a group of nodes within a larger graph that are more densely connected to each other than they are to the rest of the graph. The Leiden algorithm aims to maximize modularity by partitioning a graph into distinct, densely connected sets of nodes. Modularity is a measure to quantify the strength of the division of a graph into communities. It evaluates the density of links within communities compared to the density of links between communities. A higher modularity indicates a stronger and more defined community structure within the graph. By using a weighted mappping graph, we place more emphasis on the strongest homologies for community detection.
4. **Command generation.** For each community, we generate a complete PGGB command line in order to run it on each set of sequences separately. This eases the analysis and reduces the computational burden of building pangenome graphs.

It is important to highlight that if it is already known that the input sequences present particular rearrangements, such as rare chromosomal translocations, it may be advisable to skip the sequence partitioning and conduct the analysis with the full set of sequences.

### B.4 Validation experiments

To evaluate the accuracy and reliability of our pangenome graph construction and variant calling methods, we designed a cross-validation approach that allowed us to compare the results obtained from the graph-based method (PGGB) against those generated by the widely-used pairwise alignment tool, NUCMER, in MUMMER4 [23].

The cross-validation process begins with the extraction of FASTA sequences from the pangenome graph GFA and preparation of reference sequences. Next, variants are identified using both PGGB (with VG) and NUCMER (via a MUMMER4 script), generating a VCF file for each haplotype to ease comparison using the RealTime Genomics toolkit.

These variant files are then compared and evaluated, focusing on regions where both methods are able to call variants, an aspect that we found to be important in the HPRC cross-validation studies, wherein DIPCALL was used to find consistently-alignable regions in which comparisons were conceptually sound [18]. Finally, we collect metrics and statistics for further analysis and visualization. To simplify reproducibility, here we provide a detailed summary of the evaluation process:

1. **Extract sequences in FASTA**: The script extracts the reference paths in the GFA file and creates a new FASTA file containing these sequences.
2. **Identify variants with PGGB**: The script then identifies variants in the pangenome graph using the vg deconstruct tool with appropriate options for haplotype-based variant calling from the graph and complex allele decomposition with BiWFA and VCFLIB. The final variants are saved in a VCF format file.
3. **Pre-process the PGGB-based VCF files**: For compatibility with NUCMER, we pre-process the VCF files, normalizing alleles, removing insertions and deletions larger than 1 base pair, and removing the ALT allele if it is not present in the haplotype.
4. **Identify variants with NUCMER**: The script performs a pairwise sequence alignment between the reference and each contig in the pangenome using NUCMER. The script extracts SNPs from the NUCMER delta file using the show-snps command and generates VCF files for each aligned contig.
5. **Merge variants by haplotype**: The script then merges all VCF files for each haplotype generated by NUCMER to create a multi-haplotype VCF file.
6. **Variant evaluation**: RTG Tools’ vcfeval is used to evaluate the performance of PGGB-based variants and NUCMER-based variants by comparing true positives, false positives, and false negatives in shared callable regions. This is done for both “non-waved” and “waved” (BiWFA-normalized) PGGB-based VCF files, allowing for a direct comparison of the performance of these variant calling methods.
7. **Collect statistics**: The script computes summary statistics, such as precision, recall, and F-scores for each haplotype and writes them to TSV files. It also calculates the total number of variants called and the ratio of evaluated variants for both NUCMER and PGGB-based methods.
8. **Organize output**: Finally, the script organizes the output data, including VCF files, evaluation results, and statistics, into a specified output directory.

Although imperfect due to our lack of ground truth in the context of whole-genome alignment, this method provides a way to approximately compare the existing standard for whole-genome pairwise alignment, MUMMER4, with PGGB. We focus on SNPs and omit comparison of structural variation for diverse reasons. First, we found extracting SVs from MUMMER4 output to be problematic and poorly-supported. Second this issue remains difficult in genomics due to the multiple near-equivalent representations that a given structural variant allele may have. However, we have addressed these topics in the context of the HPRC paper [18], where significant resources were available to drive an independent evaluation of PGGB and other graph building methods. The SV validation method used in [18] involved comparing the SVs represented in the pangenome graphs (Minigraph, MC, and PGGB) to a truth set of SVs called from PacBio HiFi reads aligned to GRCh38 using three different SV discovery methods: PBSV, Sniffles with Iris, and SVIM This study found PGGB to accurately represent SVs relative to other graph construction approaches. However, SV comparison remains an ongoing challenge in the field, as multiple equivalent representations of the same variant are possible [45]. Continued work is needed to establish robust benchmarking procedures for SV calling in pangenome graphs.

### B.5 Small variants benchmark

To further evaluate the quality of genome variation captured by the pangenome graphs, we compared the variants called from the graphs to variants identified from PacBio HiFi reads by DeepVariant [46], a state-of-the-art reference-based pipeline that uses a deep neural network to call genetic variants from next-generation DNA sequencing data.

1. **Identify variants with PGGB**: The script then identifies variants in the pangenome graph using the `vg deconstruct` tool with appropriate options for haplotype-based variant calling from the graph and complex allele decomposition with BiWFA and VCFLIB. The final variants are saved in a VCF format file.
2. **Pre-process the PGGB-based VCF files**: We pre-process the VCF files, normalizing alleles, removing insertions and deletions larger than 50 base pair, and removing the ALT allele if it is not present in the haplotype.
3. **Variant evaluation**: RTG Tools’ vcfeval is used to evaluate the performance of PGGB-based variants and DeepVariant-based variants by comparing true positives, false positives, and false negatives in shared callable regions. This is done for “waved” (BiWFA-normalized) PGGB-based VCF files, allowing for a direct comparison of the performance of these variant calling methods.
4. **Collect statistics**: We compute summary statistics, such as precision, recall, and F-scores for each sample and writes them to TSV files.

Comparing small variants from the pangenome graphs to the DeepVariant reference-based sets, we observed high levels of concordance that varied, as expected, by the relative complexity of the genome (repeat content). This is true for the 3 species investigated, that is *H. sapiens* B4, *A. thaliana* B5, and tomato B6.

## Appendix C PGGB parameter settings

PGGB exposes adjustable parameters that affect the structure of the graph representing the input sequences.

Notably, modifying the mapping identity parameter (-p) significantly changes the alignment’s sensitivity. A low mapping identity increases the sensitivity, resulting in more compressed graphs. It is recommended to change this parameter depending on how divergent are the input sequences. To estimate the sequence divergence in the pangenomes, we use mash [47]. Mash is an alignment-free method that allows us to quickly derive a distance metric, the Mash distance, which estimates the mutation rate between two sequences. By calculating the Mash distance between all possible pairs of the input sequences, we can obtain the maximum distance, and therefore an estimation of the divergence in the pangenome. We recommend specifying a mapping identity threshold close to 100 *max_distance*%. For example, if the input genomes have a maximum divergence of 2%, we can run PGGB by specifying a mapping identity threshold of 100 − 2% = 98%. However, it should be noted that if users are interested in modeling even older homologies in the final graph, a lower value than the suggested one may be specified.

The segment length parameter (-s) determines the minimum sequence length for the initial mapping step performed by WFMASH. This parameter works in conjunction with the mapping identity (-p) to establish the structure of the final pangenome graph. When WFMASH performs the mapping step, it only considers segments that meet the specified segment length and have an approximate identity greater than or equal to the mapping identity minimum. The choice of segment length depends on the characteristics of the sequences being used and the presence of repeats in the pangenome. For smaller pangenome graphs or those with few repeats, a lower segment length can be used. By default, we use a segment length of 5000 bps. For larger contexts or those with many repeats, setting a higher segment length helps in building not-too-complex graphs. By setting a long segment length, the final graph will have a more linear structure, representing long collinear regions of the input sequences. As a general guideline, the segment length should be larger than the size of common repeats in the pangenome, such as transposons.

**Fig. B4:**
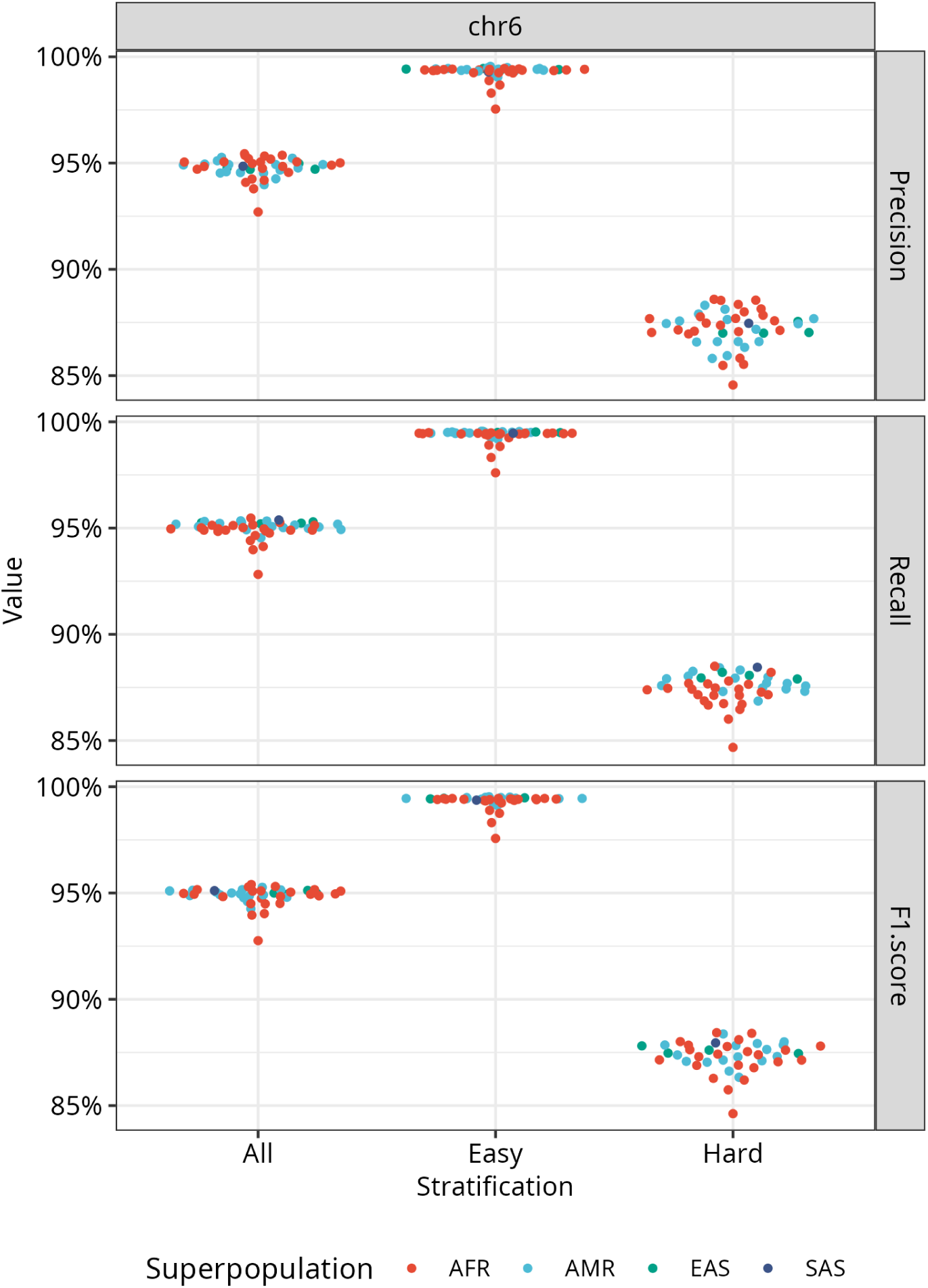
Precision, recall, and F1-score of small variants in the *H. sapiens* chromosome 6 pangenome graph relative to HiFi–DeepVariant calls. Comparisons are made whole-chromosome and then stratified by the GIAB (v.3.0) genomic context. The 44 samples evaluated are colored by superpopulation. AFR = African, AMR = Ad Mixed American, EAS = East Asian, SAS = South Asian.

**Fig. B5:**
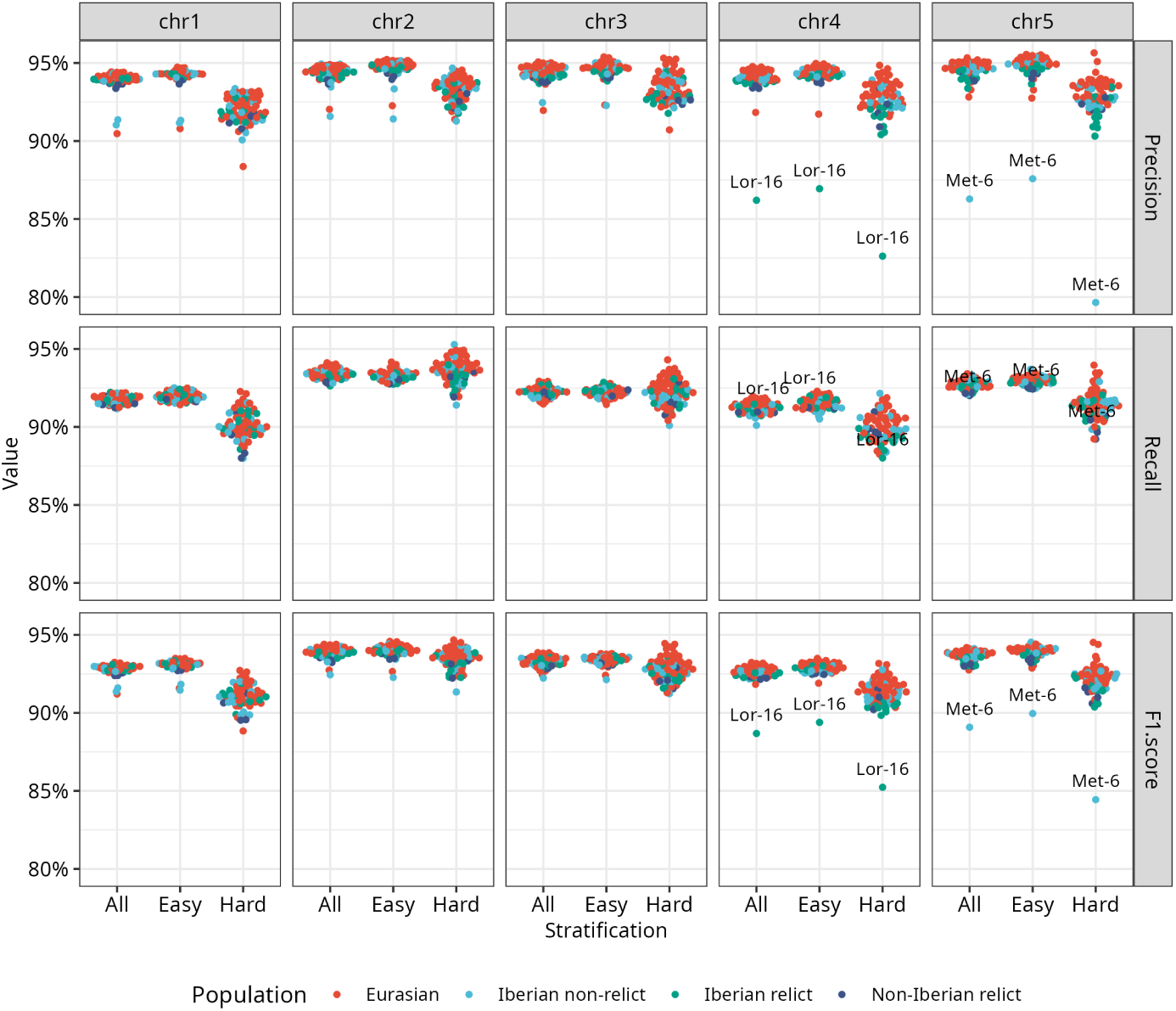
Precision, recall, and F1-score of small variants in the *A. thaliana* pangenome graph relative to HiFi–DeepVariant calls. Comparisons are made whole-genome and then stratified by genomic context. Easy and Hard regions exclude and include, respectively, rDNA, centromere, and Trasposable Elements. The 64 samples evaluated are colored by population. The low precision for Lor-16 and Met-6 is due to the high heterozygosity of these 2 samples.

The minimum match length parameter (-k) determines the minimum length that matches in the WFMASH alignment must have to avoid being filtered out by SEQWISH for graph construction. Filtering short matches reduces graph complexity while creating locally under-aligned regions in the graph that are subsequently fixed by SMOOTHXG. By default, alignment matches shorter than 23 bps are discarded. Higher values, such as 47 bps or 79 bps, help reduce graph complexity in difficult regions of the genomes, like alpha-satellites and centromeres. For more divergent genomes, it is advisable to not increase these values too much. In such cases, genomes may not share long alignment matches due to the accumulation of multiple mutations over time. Then, setting a too-high minimum match length can result in the loss of the majority of the alignments and affect the representation of the relationships between the genomes in the final graph.

An explanation of other PGGB parameters can be found online at https://pggb.readthedocs.io/.

## Appendix D Phylogeny

We consider the nodes of pangenome variation graphs as alleles, and work with simple counts of these alleles in genomes. This approach projects the pangenome into a matrix form that can readily be utilized in diverse, and standard, downstream population genetics applications (D7). Compared to traditional SNP-based trees using the S288C reference, our method more accurately clusters related haplotypes, specifically affecting CKB, AIF, and BPK strains. The clustering observed in PGGB’s data reflects the closely related evolutionary trajectories of the two haplotypes in a phased diploid genome [39]; e.g. CKB-h1 and CKB-h2, respectively refer, to the first and the second haplotype of the CKB strain.

## Appendix E Comparison with Minigraph-Cactus

Minigraph-Cactus (MC) relies on Minigraph’s graph [7], which only includes variations of at least 50 bps, and applies the Cactus aligner [48] on those graphs to produce base-level pangenome graphs. MC relies on choosing a reference sample that anchors the entire graph and is treated differently than the other samples. In particular, the reference sample is never clipped. We applied MC (version 2.7.0) to build graphs for all pangenomes in this study. MC’s graphs exclude, on average, 1.8% (in *H. sapiens* chromosome 6) to 22.1% (in *E. coli*) of the sequence of the pangenome investigated E8. For the primate pangenome, MC built a graph that resembles the genome used to anchor the graph during its construction (GRCh38), while the sequence clipping removes all centromeres, a big part of gibbons’s haplotypes, and leads to a different graph-based phylogenetic tree E9. For the *A. thaliana* pangenome, MC clipped both centromeric and not centromeric sequence (E10a), with the not centromeric regions containing several kinds of genes (E10b).

## Appendix F Pangenome openness

We used PANACUS [49] to calculate the coverage and pangenome growth curve (with estimated growth parameters) for base pairs with quorum thresholds 0, 1, 0.5, and 0.1. The *E. coli* pangenome growth curve, made with 500 genomes, is far from showing signs of saturation, with each new genome adding a significant amount of new sequence in the pangenome (F11). On the contrary, the tomato pangenome is relatively much more closed, with a growth curve close to saturation (F12). The graphs thus recover coarse, species-level evolutionary parameters which we anticipate based on prior studies [50, 51].

## Appendix G PGGB and NUCMER alignment differences

The *athaliana82* pangenome displays the lowest F1-score (0.920053) when comparing SNVs called by PGGB to those called by NUCMER. This is primarily due to a high number of false negative SNVs, that is SNVs that are called by NUCMER but not by PGGB. As shown in G14, the *athaliana82* pangenome has the highest density of very closely spaced SNVs, with many SNVs located less than 2 bp apart. PGGB (which relies on WFMASH alignments) and NUCMER handle these regions differently due to the underlying alignments they use. In particular, NUCMER allows for more mismatches in its alignments, leading to calling a higher number of close SNVs compared to PGGB. G15 provides an illustrative example using IGV, showing a region where the NUCMER alignment contains several close SNVs, while the WFMASH alignment (used by PGGB) represents the variation differently, resulting in fewer called SNVs. This issue also affects the other analyzed pangenomes, but to a much lesser degree. The *athaliana82* pangenome, with its particularly high density of nearby SNVs, is the most impacted, resulting in the lowest overall concordance between PGGB and NUCMER SNV calls.

## Appendix H Random graph model for mapping sparsification

PGGB can employ a random sparsification approach to reduce the computational complexity of all-vs-all pairwise alignments. This is particularly important for large datasets, where the number of pairwise comparisons grows quadratically with the number of genomes.

Although this sparsification can reduce sensitivity, when large numbers of homologous genomes from the same or related species are aligned, transitive recovery of pairwise relationships due to the graph induction step ensures that even if we retain only a fraction of the total mappings, all genomes will still be aligned to each other in all homologous regions.

The sparsification procedure is applied to the mapping graph, which represents the pairwise mappings between all genomes in the dataset. Each node in the mapping graph represents a genome (or haploid copy), and edges represent mappings between these genomes.

When reasoning about the sparsification process, we focus on a local region of the mapping graph corresponding to a homologous region between different genomes. Let’s define the key components within this local region:

- *N* : number of nodes in the local region of the mapping graph (representing haploid genome copies)
- *E*: number of edges in the local region of the mapping graph (representing mappings between genomes in the homologous region)

When our input alignment is completely all-to-all, the graph would be complete, with 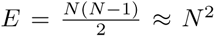. This graph will be fully connected, and when we build a pangenome graph from this region, we would expect it to contain a single component with all genomes due to walks over a minimal set of nodes. Although mappings are generally very cheap to compute, it is very expensive to compute the entire set of alignments. Thus, we sparsify the mapping graph by randomly removing pairwise mappings. This reduces the amount of pairwise alignment that must be done to compute the input for graph induction in seqwish.

We must determine how many edges we can remove in the mapping graph without disrupting its connectivity. If the mapping graph becomes unconnected, homologous regions will not align to each other in the pangenome graph due to mapping sparsification. To determine a suitable threshold for connectivity, we build on the Erdős–Rényi random graph model. The Erdős–Rényi model predicts that as *N* approaches infinity, a random graph with *N* nodes is almost certainly fully connected as long as pairs of nodes are connected with a probability 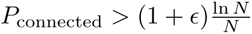, where *ɛ* is a small constant.

We set a sparsification parameter that uses a hash of each mapping record to drop mappings with a probability *P*_sparse_≫ *P*_connected_. This ensures that while some edges are removed, the giant component encompassing critical homologous relationships is preserved with high probability.

Specifically, PGGB uses the following heuristic based on the Erdős–Rényi model to set the sparse mapping fraction (*n* is the number of haplotypes):

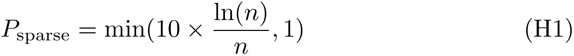

This allows PGGB to reconstruct all transitive relationships in the variation graph without needing to directly compute all pairwise alignments, avoiding the expected *O*(*N* ^2^) costs that would occur if *P*_sparse_ = 1. This dramatically reduces the runtime of alignment and graph induction with negligible effect on accuracy.

The choice of the sparsification factor balances the trade-off between graph completeness and computational efficiency. PGGB uses a default value that has been empirically observed to work well across a range of datasets. This random sparsification approach is unbiased, as it does not rely on any prior knowledge of the relationships between genomes. It provides a principled way to reduce computational complexity while preserving key homology information in the pangenome graph construction process.

We anticipate that further improvements to this sparsification procedure may be yielded through the application of neighbor joining trees over sequence similarity metrics in conjunction with randomly sampled alignments from the complete mapping graph.

**Fig. B6:**
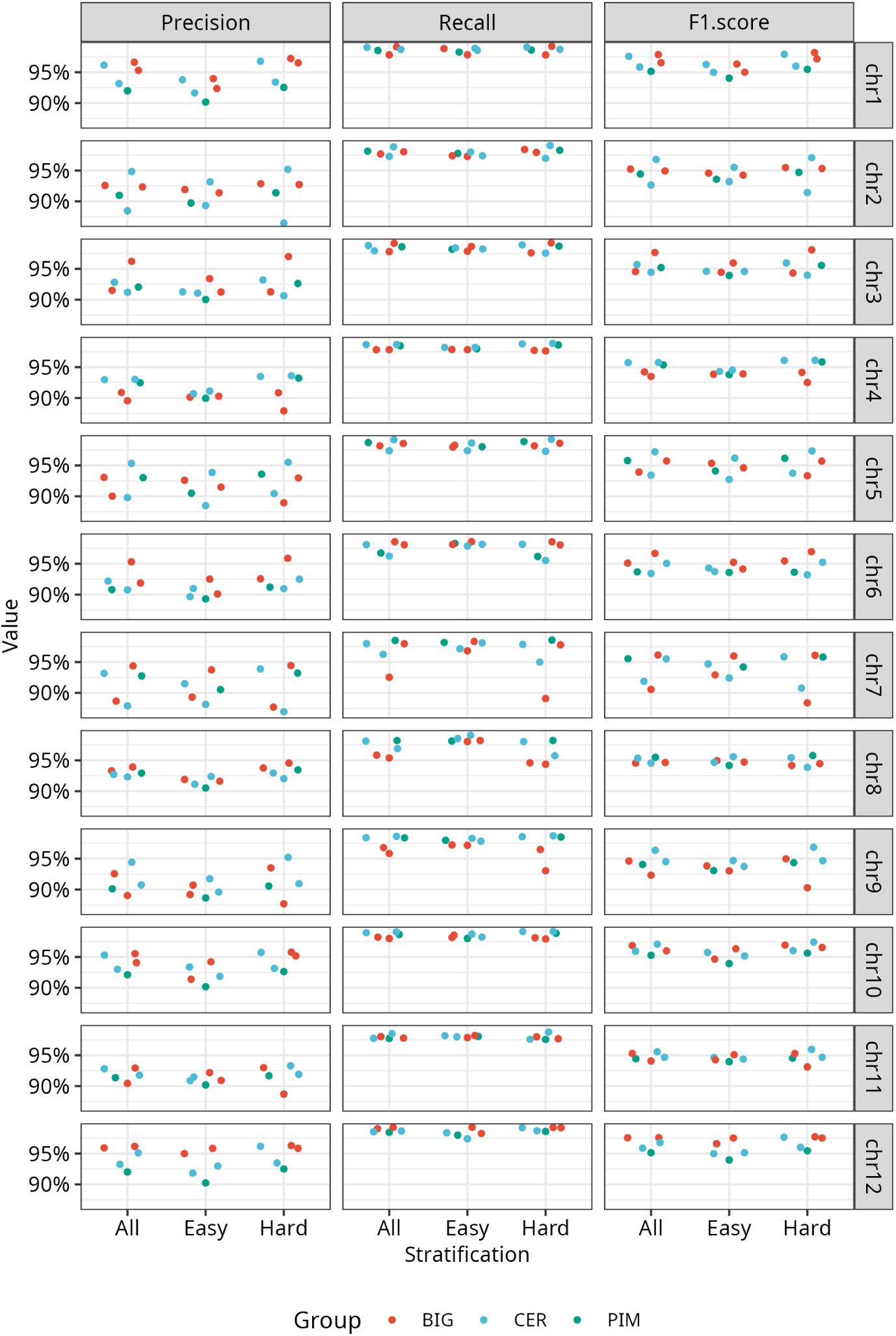
Precision, recall, and F1-score of small variants in the tomato pangenome graph relative to HiFi–DeepVariant calls. Comparisons are made whole-genome and then stratified by genomic context. Easy and Hard regions exclude and include, respectively, Transposable elements. The 5 samples evaluated are colored by group. BIG = *S. lycopersicum*, big-fruited tomato; CER = *S. lycopersicum var. cerasiforme*, cherry tomato; PIM = *S. pimpinellifolium*, the progenitor of cultivated tomatoes.

**Fig. D7:**
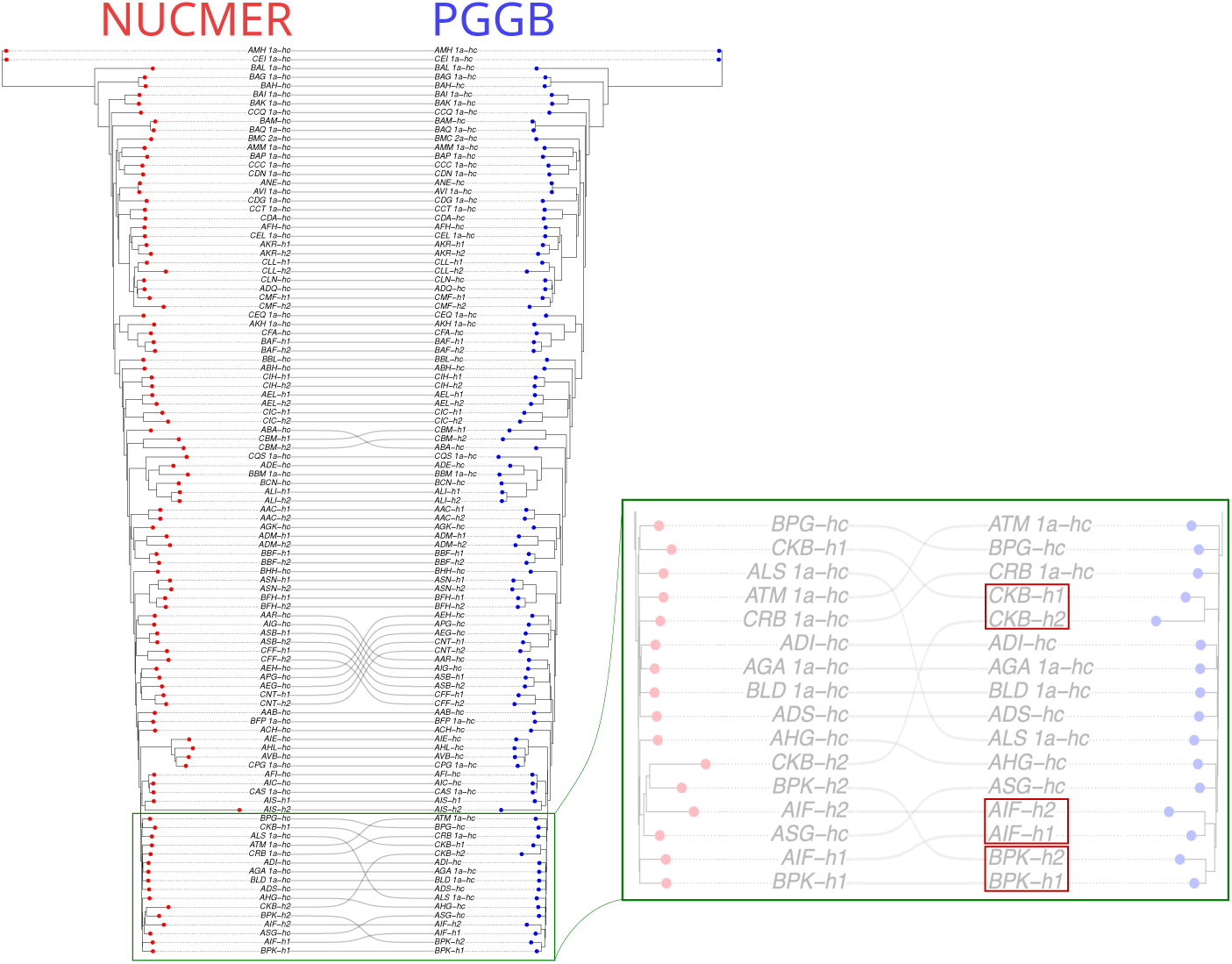
Pangenomic phylogeny in yeast. We compared the phylogenetic trees of 142 yeast genomes: one created using single nucleotide variants (SNVs) via MUMMER (”NUCMER”) and another generated from graph data using node-coverage vectors (”PGGB”). Both trees largely align, but the PGGB-based tree correctly groups haplotypes from several diploid assemblies (in red).

**Fig. E8:**
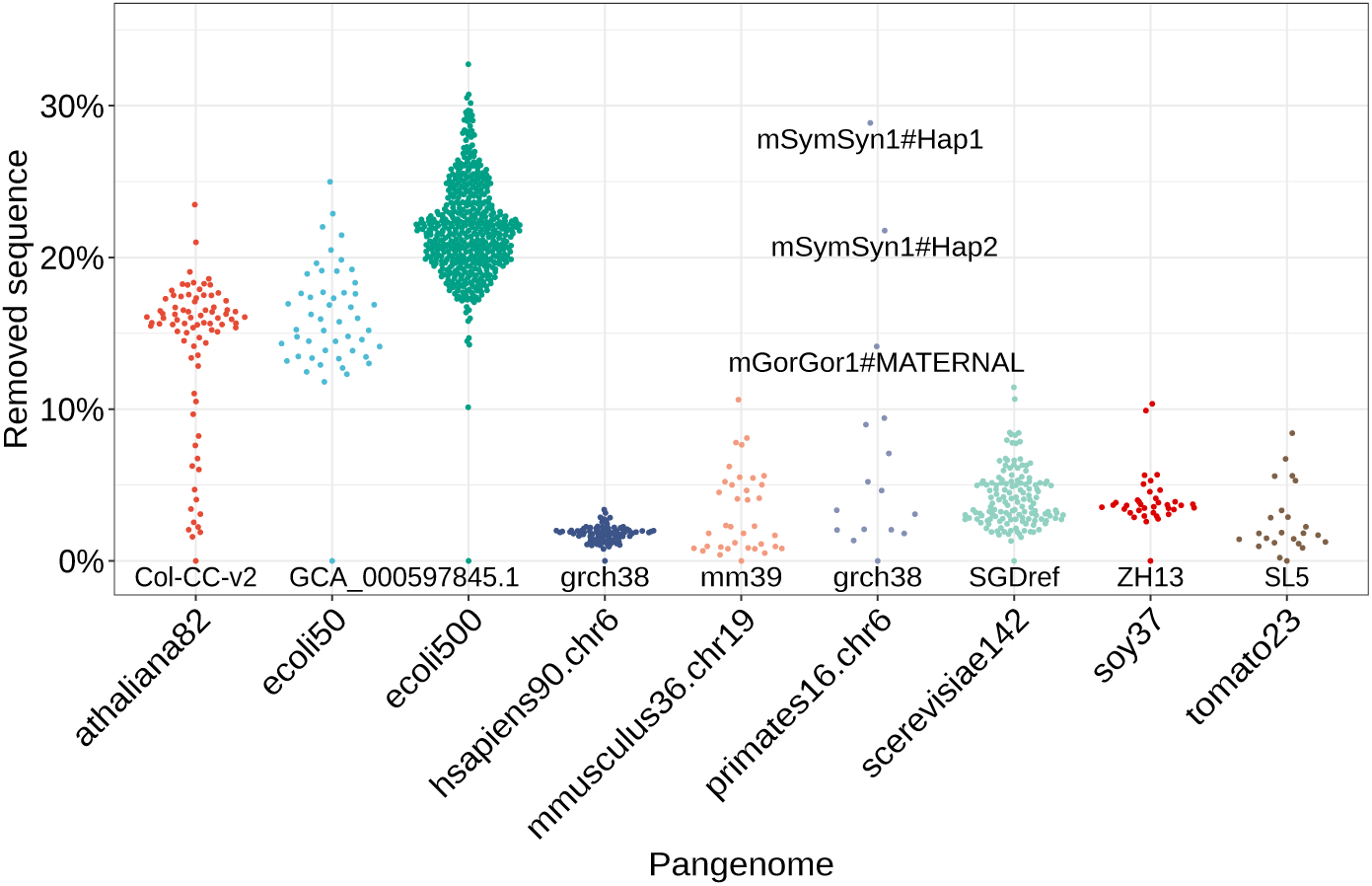
Removed pangenome sequence in Minigraph-Cactus’ graphs. Labels for reference samples and a few outliers are reported. In the *E. coli* pangenome with 500 samples, two extreme cases are present (not shown) which could be misassemblies: GCA 902141745.2 (VRES-hospital6495300 strain, 170 kbps, flagged by GenBank as too small and with unverified source organism) was completely clipped, while GCA 025790905.1 (2020CK-00232 strain, 5.6 Mbps, flagged by GenBank as contaminated and with unverified source organism) lost 92.78% of its sequence. In the primates graph, gibbon’s haplotypes (mSymSyn) lost more than 20% of their sequence.

**Fig. E9:**
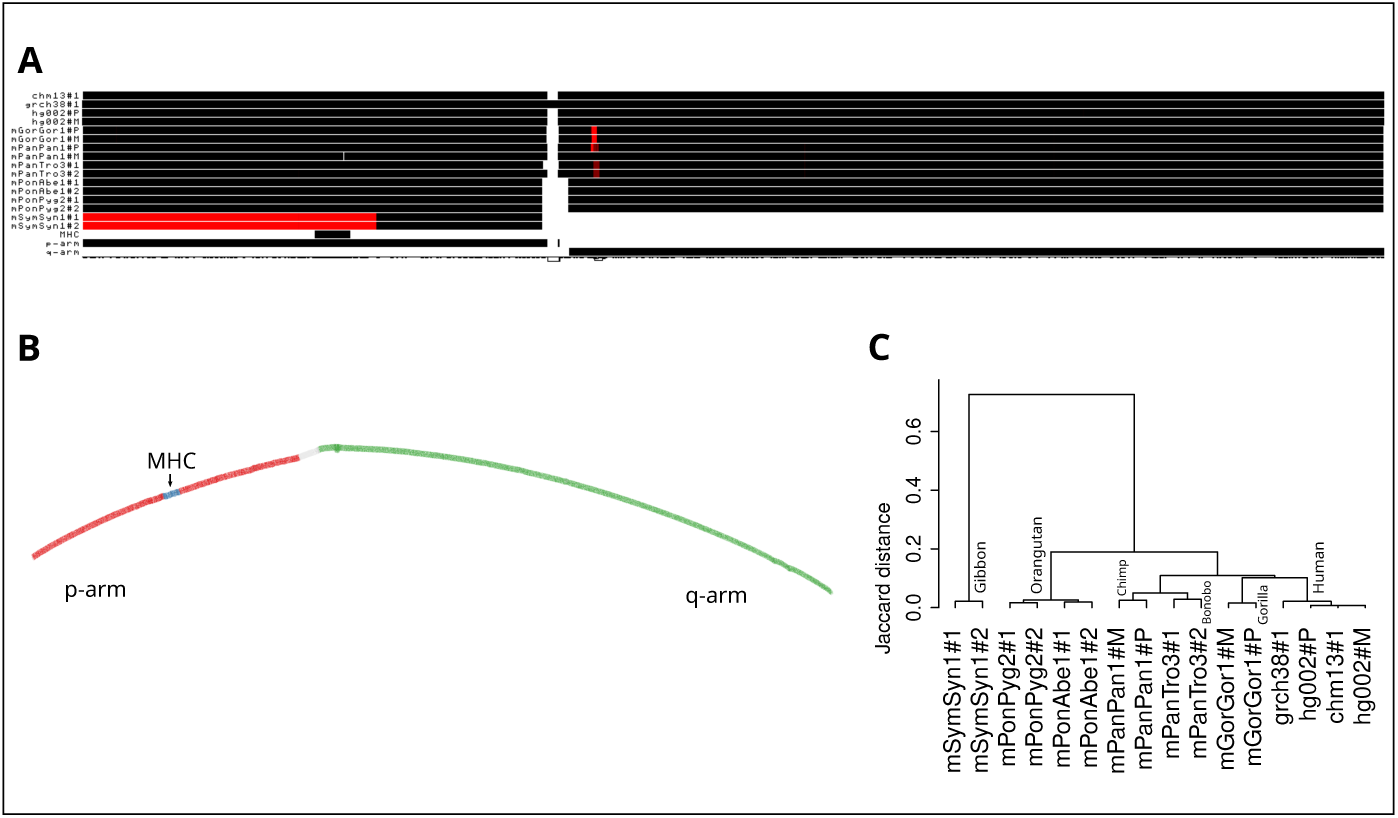
Primate pangenome graph with Minigraph-Cactus. (A) 1D pangenome graph visualization using 16 haplotype-resolved primate assemblies homologous to human chromosome 6. T2T-CHM13 annotations (Major Histocompatibility Complex, p-arm, q-arm) are shown. Black and red indicate regions, respectively, in forward and reverse with respect to the pangenome sequence (the sequence obtained by concatenating all graph nodes). The p-arm region with the MHC is inverted in Gibbon. Centromeric regions appear all clipped, except for GRCh38, as it was used by MC to anchor the pangenome graph. (B) 2D visualization, rendered with the same human chromosomal annotations in GFAESTUS [15]. (C) Using ODGI [6], we extract a pairwise distance matrix based on in-graph Jaccard metrics over shared base pairs. This distance matrix yields a phylogenetic tree that does not match previous results based on SNPs [16] and obtained with the PGGB graph, probably due to MC’s sequence clipping.

**Fig. E10:**
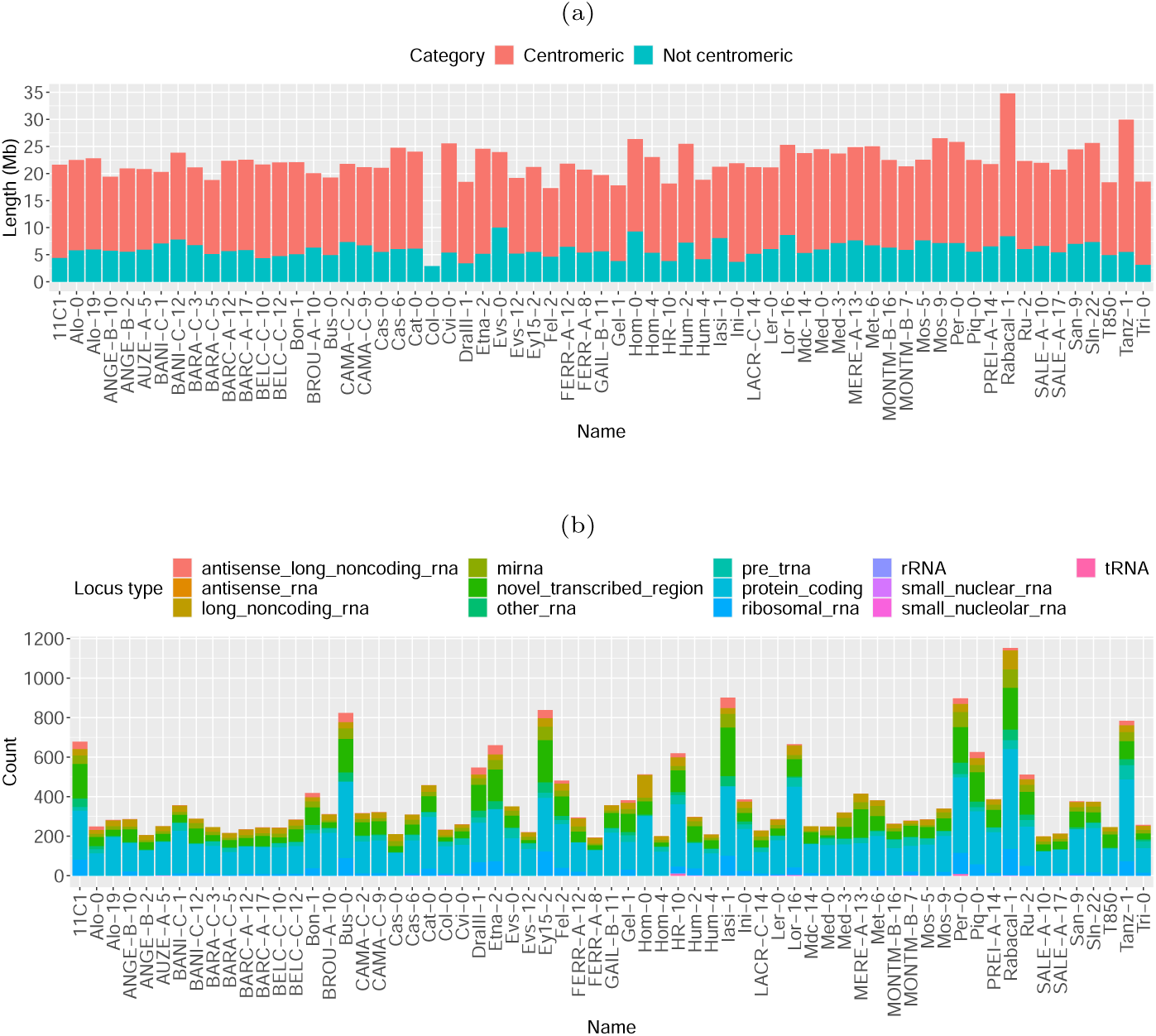
Minigraph-Cactus removes centromeric and gene-containing not centromeric regions in *A.thaliana*. The graph was made with 82 genomes, using Col-CC-v2 (GCA 028009825.2) as a reference to anchor the graph, and annotating the genomes assembled from HiFi reads (65 in total). a) Amount of sequence clipped by Minigraph-Cactus, classified as centromeric or not centromeric. b) Loci classification in the clipped not centromeric sequences.

**Fig. F11:**
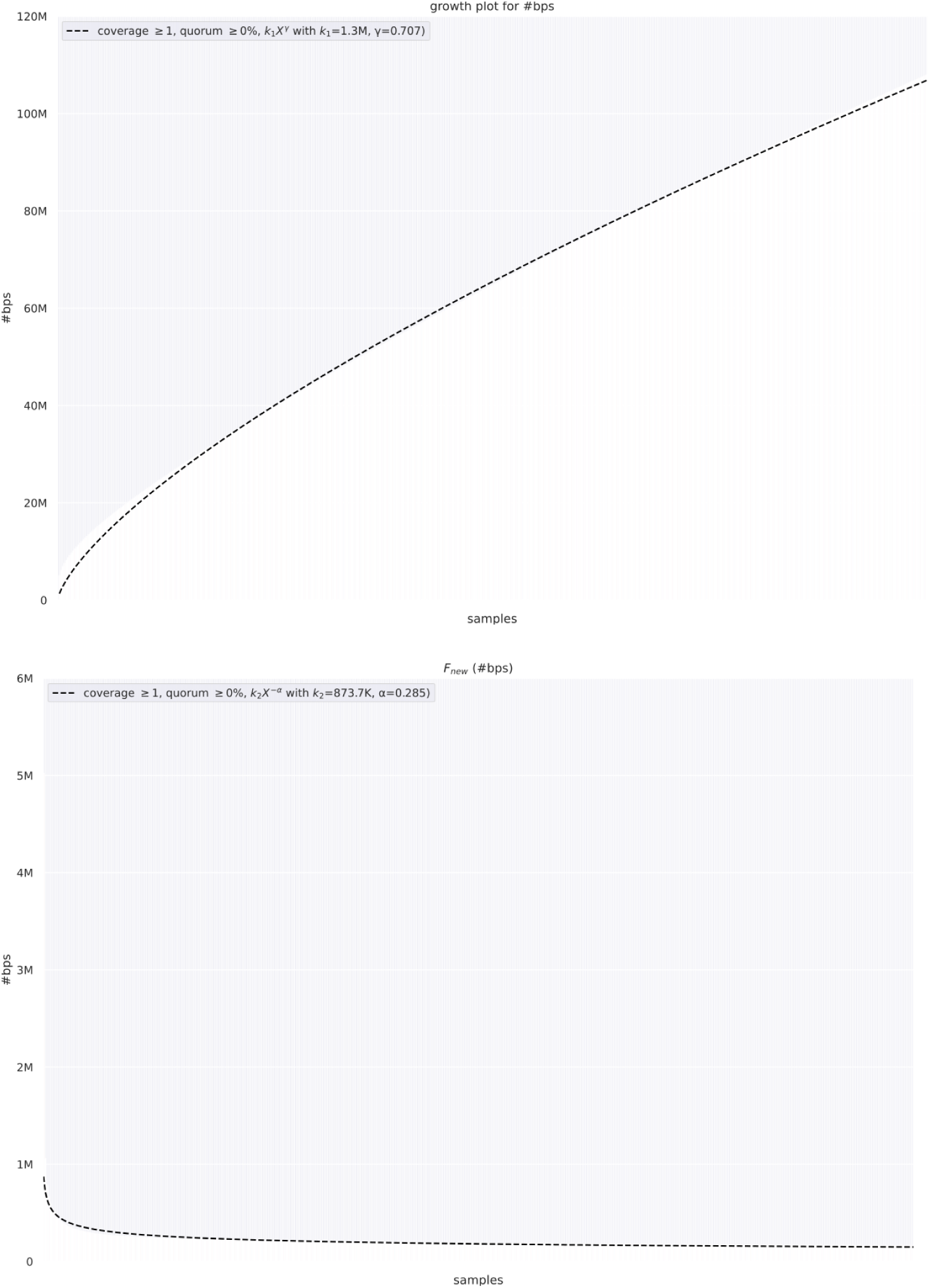
Pangenome growth curve with estimated growth parameters and coverage histogram for the *E. coli* pangenome made with 500 samples. Sample ranks on the x-axis are omitted. Only the fitted curves are visualized.

**Fig. F12:**
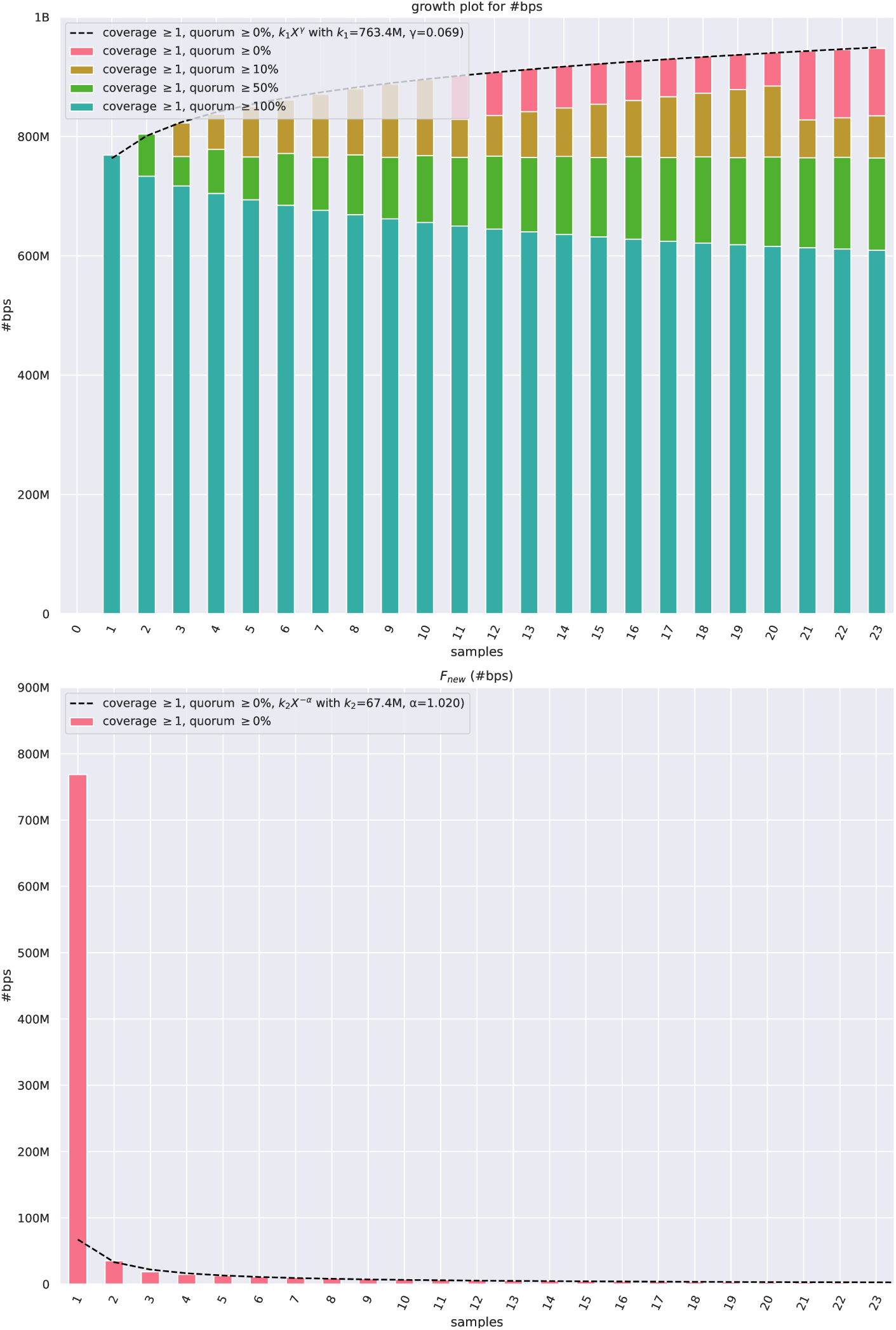
Pangenome growth curve with estimated growth parameters and coverage histogram for the tomato pangenome made with 23 samples.

**Fig. F13:**
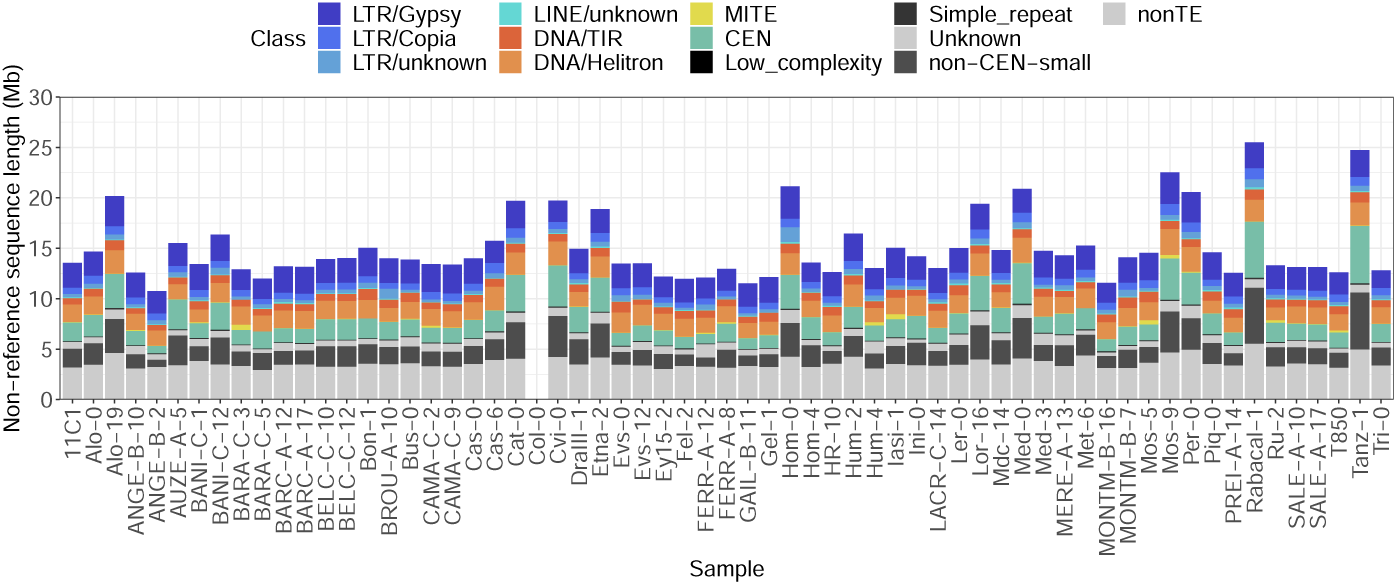
Non-reference sequence annotation in the *A. thaliana* pangenome. The PGGB pangenome graph was made with 82 genomes, annotating the ones assembled from HiFi reads (63 out of 65, as we were unable to annotate IP-San-9 and IP-Sln-22 because of a bug in the Extensive de novo TE Annotator [52]). We used Col-CC-v2 (GCA 028009825.2, proposed as the next version of *Arabidopsis thaliana* reference) as a reference to define the non-reference sequences) as a reference to define the non-reference sequences. Col-CC is the Col-0 Community Consensus assembly, which was generated by integrating high-quality genome assemblies of 13 different Col-0 accessions from different groups worldwide using HiFi reads. These 13 Col-0 accessions include GCA 946499705.1, which is part of the *A. thaliana* pangenome we considered. The consensus approach used to create Col-CC-v2 explains the low amount of non-reference sequences observed for the Col-0 assembly included in our analysis (that is, GCA 946499705.1).

**Fig. G14:**
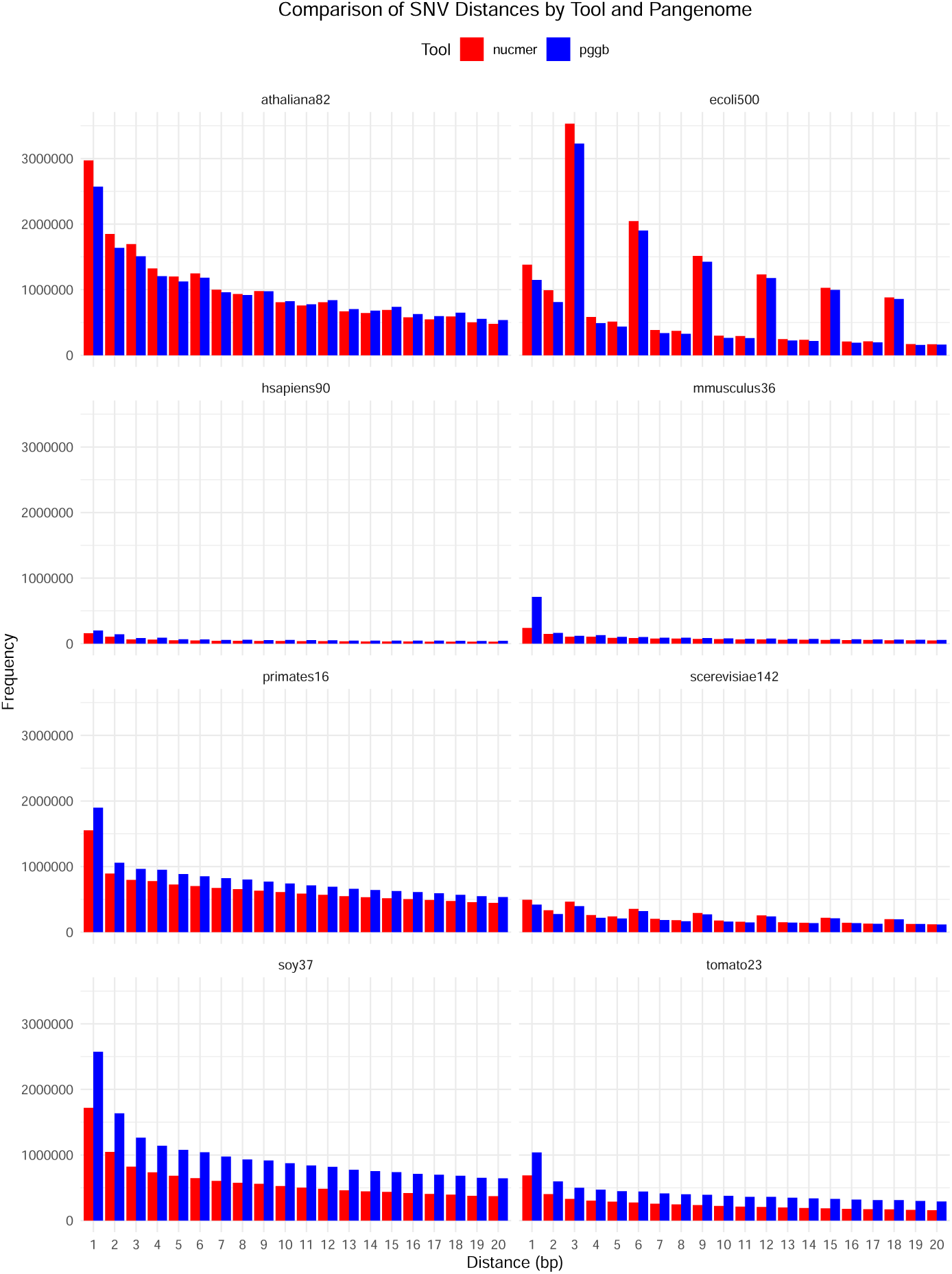
Pangenome SNV distances. For PGGB-called SNVs and NUCMER-called SNVS, we show the distribution of their distances, zooming in the interval 0-20. *athaliana82* display the highest number of closely spaced SNVs.

**Fig. G15:**
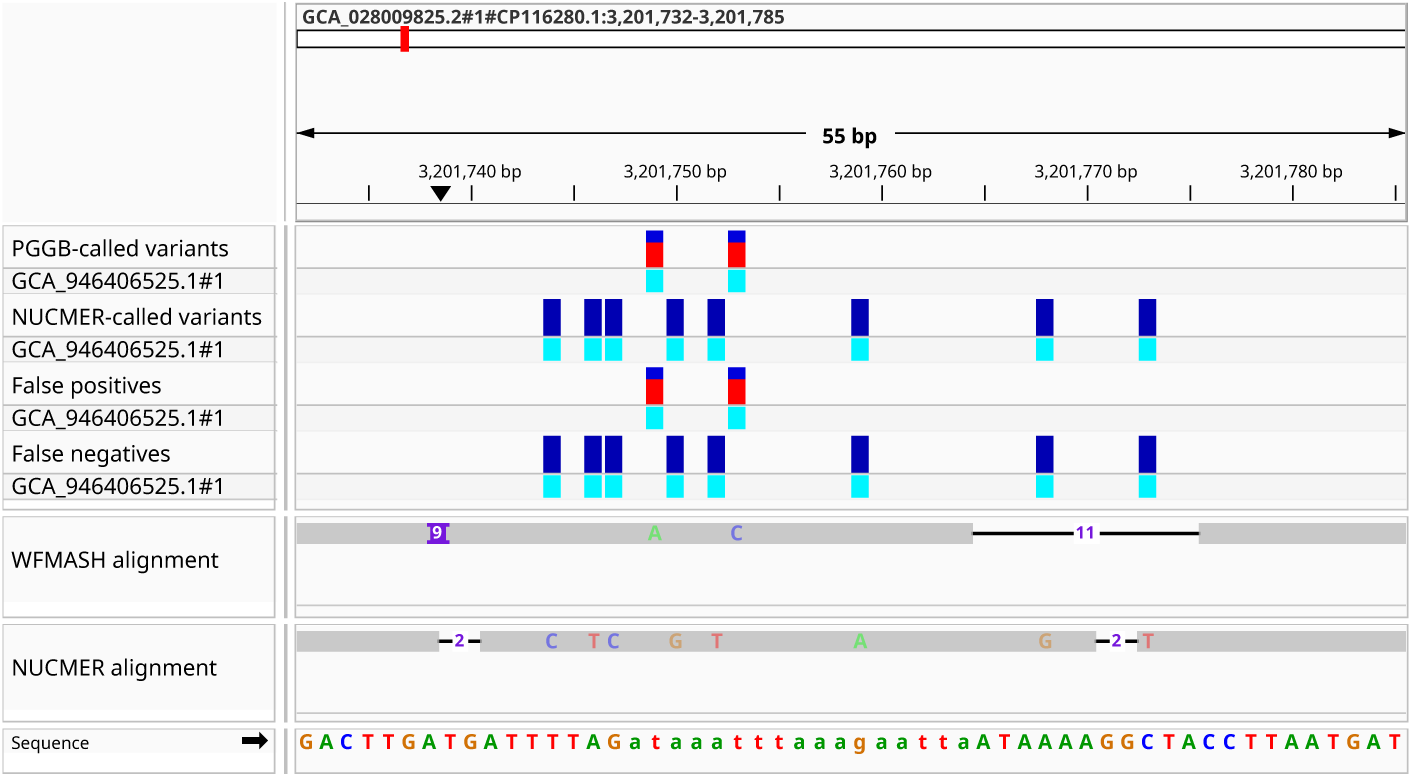
NUCMER allows for more mismatches in SNV clusters. Integrative Genomics Viewer (IGV) visualization ([53] of the alignment between 2 *A. thaliana* accessions, GCA 946406525.1 against GCA 028009825.2. Variants are expressed using GCA 028009825.2 as the reference genome. The figure shows a region on chromosome 1, GCA 028009825.2#1#CP116280.1:3,201,732-3,201,785. From top to bottom, the tracks show the SNVs called by PGGB, SNVs called by NUCMER, false positives (SNVs called by PGGB but not by NUCMER), false negatives (SNVs called by NUCMER but not by PGGB), WFMASH alignments (on which PGGB relies) and NUCMER alignment.

## References

[1] Garrison, E., Sirén, J., Novak, A.M., Hickey, G., Eizenga, J.M., Dawson, E.T., Jones, W., Garg, S., Markello, C., Lin, M.F., Paten, B., Durbin, R.: Variation graph toolkit improves read mapping by representing genetic variation in the reference. Nature Biotechnology 36(9), 875–879 (2018). 10.1038/nbt.4227

[2] Paten, B., Novak, A.M., Eizenga, J.M., Garrison, E.: Genome graphs and the evolution of genome inference. Genome Res. 27(5), 665–676 (2017). 10.1101/gr.214155.116

[3] Garrison, E.: Graphical pangenomics (2019). 10.17863/CAM.41621

[4] Eizenga, J.M., Novak, A.M., Sibbesen, J.A., Heumos, S., Ghaffaari, A., Hickey, G., Chang, X., Seaman, J.D., Rounthwaite, R., Ebler, J., Rautiainen, M., Garg, S., Paten, B., Marschall, T., Sirén, J., Garrison, E.: Pangenome graphs. Annual Review of Genomics and Human Genetics 21(1), 139–162 (2020). 10.1146/annurev-genom-120219-080406

[5] Armstrong, J., Hickey, G., Diekhans, M., Fiddes, I.T., Novak, A.M., Deran, A., Fang, Q., Xie, D., Feng, S., Stiller, J., Genereux, D., Johnson, J., Marinescu, V.D., Alföldi, J., Harris, R.S., Lindblad-Toh, K., Haussler, D., Karlsson, E., Jarvis, E.D., Zhang, G., Paten, B.: Progressive cactus is a multiple-genome aligner for the thousand-genome era. Nature 587(7833), 246–251 (2020). 10.1038/s41586-020-2871-y

[6] Guarracino, A., Heumos, S., Nahnsen, S., Prins, P., Garrison, E.: ODGI: understanding pangenome graphs. Bioinformatics (2022). 10.1093/bioinformatics/btac308

[7] Li, H., Feng, X., Chu, C.: The design and construction of reference pangenome graphs with minigraph. Genome Biology 21(1) (2020). 10.1186/s13059-020-02168-z

[8] Hickey, G., Monlong, J., Ebler, J., Novak, A.M., Eizenga, J.M., Gao, Y., Abel, H.J., Antonacci-Fulton, L.L., Asri, M., Baid, G., Baker, C.A., Belyaeva, A., Billis, K., Bourque, G., Buonaiuto, S., Carroll, A., Chaisson, M.J.P., Chang, P.-C., Chang, X.H., Cheng, H., Chu, J., Cody, S., Colonna, V., Cook, D.E., Cook-Deegan, R.M., Cornejo, O.E., Diekhans, M., Doerr, D., Ebert, P., Ebler, J., Eichler, E.E., Fairley, S., Fedrigo, O., Felsenfeld, A.L., Feng, X., Fischer, C., Flicek, P., Formenti, G., Frankish, A., Fulton, R.S., Garg, S., Garrison, E., Garrison, N.A., Giron, C.G., Green, R.E., Groza, C., Guarracino, A., Haggerty, L., Hall, I.M., Harvey, W.T., Haukness, M., Haussler, D., Heumos, S., Hoekzema, K., Hourlier, T., Howe, K., Jain, M., Jarvis, E.D., Ji, H.P., Kenny, E.E., Koenig, B.A., Kolesnikov, A., Korbel, J.O., Kordosky, J., Koren, S., Lee, H., Lewis, A.P., Liao, W.-W., Lu, S., Lu, T.-Y., Lucas, J.K., Magalhães, H., Marco-Sola, S., Marijon, P., Markello, C., Marschall, T., Martin, F.J., McCartney, A., McDaniel, J., Miga, K.H., Mitchell, M.W., Mountcastle, J., Munson, K.M., Mwaniki, M.N., Nattestad, M., Nurk, S., Olsen, H.E., Olson, N.D., Pesout, T., Phillippy, A.M., Popejoy, A.B., Porubsky, D., Prins, P., Puiu, D., Rautiainen, M., Regier, A.A., Rhie, A., Sacco, S., Sanders, A.D., Schneider, V.A., Schultz, B.I., Shafin, K., Sibbesen, J.A., Sirén, J., Smith, M.W., Sofia, H.J., Tayoun, A.N.A., Thibaud-Nissen, F., Tomlinson, C., Tricomi, F.F., Villani, F., Vollger, M.R., Wagner, J., Walenz, B., Wang, T., Wood, J.M.D., Zimin, A.V., Zook, J.M., Marschall, T., Li, H., Paten, B.: Pangenome graph construction from genome alignments with minigraph-cactus. Nat Biotechnol (2023). 10.1038/s41587-023-01793-w

[9] Noll, N., Molari, M., Shaw, L.P., Neher, R.A.: PanGraph: scalable bacterial pan-genome graph construction (2022). 10.1101/2022.02.24.481757

[10] Garrison, E., Guarracino, A.: Unbiased pangenome graphs. Bioinformatics 39(1) (2022). 10.1093/bioinformatics/btac743

[11] Minkin, I., Pham, S., Medvedev, P.: TwoPaCo: an efficient algorithm to build the compacted de bruijn graph from many complete genomes. Bioinformatics 33(24), 4024–4032 (2016). 10.1093/bioinformatics/btw609

[12] Chin, C.-S., Behera, S., Khalak, A., Sedlazeck, F.J., Sudmant, P.H., Wagner, J., Zook, J.M.: Multiscale analysis of pangenomes enables improved representation of genomic diversity for repetitive and clinically relevant genes. Nat Methods (2023). 10.1038/s41592-023-01914-y

[13] Sullivan, P.F., Meadows, J.R.S., Gazal, S., Phan, B.N., Li, X., Genereux, D.P., Dong, M.X., Bianchi, M., Andrews, G., Sakthikumar, S., Nordin, J., Roy, A., Christmas, M.J., Marinescu, V.D., Wang, C., Wallerman, O., Xue, J., Yao, S., Sun, Q., Szatkiewicz, J., Wen, J., Huckins, L.M., Lawler, A., Keough, K.C., Zheng, Z., Zeng, J., Wray, N.R., Li, Y., Johnson, J., Chen, J., Paten, B., Reilly, S.K., Hughes, G.M., Weng, Z., Pollard, K.S., Pfenning, A.R., Forsberg-Nilsson, K., Karlsson, E.K., Lindblad-Toh, K., Andrews, G., Armstrong, J.C., Bianchi, M., Birren, B.W., Bredemeyer, K.R., Breit, A.M., Christmas, M.J., Clawson, H., Damas, J., Di Palma, F., Diekhans, M., Dong, M.X., Eizirik, E., Fan, K., Fanter, C., Foley, N.M., Forsberg-Nilsson, K., Garcia, C.J., Gatesy, J., Gazal, S., Genereux, D.P., Goodman, L., Grimshaw, J., Halsey, M.K., Harris, A.J., Hickey, G., Hiller, M., Hindle, A.G., Hubley, R.M., Hughes, G.M., Johnson, J., Juan, D., Kaplow, I.M., Karlsson, E.K., Keough, K.C., Kirilenko, B., Koepfli, K.-P., Korstian, J.M., Kowalczyk, A., Kozyrev, S.V., Lawler, A.J., Lawless, C., Lehmann, T., Levesque, D.L., Lewin, H.A., Li, X., Lind, A., Lindblad-Toh, K., Mackay-Smith, A., Marinescu, V.D., Marques-Bonet, T., Mason, V.C., Meadows, J.R.S., Meyer, W.K., Moore, J.E., Moreira, L.R., Moreno-Santillan, D.D., Morrill, K.M., Muntané, G., Murphy, W.J., Navarro, A., Nweeia, M., Ortmann, S., Osmanski, A., Paten, B., Paulat, N.S., Pfenning, A.R., Phan, B.N., Pollard, K.S., Pratt, H.E., Ray, D.A., Reilly, S.K., Rosen, J.R., Ruf, I., Ryan, L., Ryder, O.A., Sabeti, P.C., Schäffer, D.E., Serres, A., Shapiro, B., Smit, A.F.A., Springer, M., Srinivasan, C., Steiner, C., Storer, J.M., Sullivan, K.A.M., Sullivan, P.F., Sundström, E., Supple, M.A., Swofford, R., Talbot, J.-E., Teeling, E., Turner-Maier, J., Valenzuela, A., Wagner, F., Wallerman, O., Wang, C., Wang, J., Weng, Z., Wilder, A.P., Wirthlin, M.E., Xue, J.R., Zhang, X.: Leveraging base-pair mammalian constraint to understand genetic variation and human disease. Science 380(6643) (2023). 10.1126/science.abn2937

[14] Doerr, D., Marijon, P., Marschall, T.: GFAffix identifies walk-preserving shared affixes in variation graphs and collapses them into a non-redundant graph structure (2023). https://github.com/marschall-lab/GFAffix

[15] Fischer, C., Garrison, E.: chfi/gfaestus: a pangenome graph browser. Zenodo (2022). 10.5281/ZENODO.6954035. https://zenodo.org/record/6954035

[16] Cagan, A., Theunert, C., Laayouni, H., Santpere, G., Pybus, M., Casals, F., Prüfer, K., Navarro, A., Marques-Bonet, T., Bertranpetit, J., Andrés, A.M.: Natural selection in the great apes. Mol Biol Evol 33(12), 3268– 3283 (2016). 10.1093/molbev/msw215

[17] Logsdon, G.A., Vollger, M.R., Hsieh, P., Mao, Y., Liskovykh, M.A., Koren, S., Nurk, S., Mercuri, L., Dishuck, P.C., Rhie, A., de Lima, L.G., Dvorkina, T., Porubsky, D., Harvey, W.T., Mikheenko, A., Bzikadze, A.V., Kremitzki, M., Graves-Lindsay, T.A., Jain, C., Hoekzema, K., Murali, S.C., Munson, K.M., Baker, C., Sorensen, M., Lewis, A.M., Surti, U., Gerton, J.L., Larionov, V., Ventura, M., Miga, K.H., Phillippy, A.M., Eichler, E.E.: The structure, function and evolution of a complete human chromosome 8. Nature 593(7857), 101–107 (2021). 10.1038/s41586-021-03420-7

[18] Liao, W.-W., Asri, M., Ebler, J., Doerr, D., Haukness, M., Hickey, G., Lu, S., Lucas, J.K., Monlong, J., Abel, H.J., Buonaiuto, S., Chang, X.H., Cheng, H., Chu, J., Colonna, V., Eizenga, J.M., Feng, X., Fischer, C., Fulton, R.S., Garg, S., Groza, C., Guarracino, A., Harvey, W.T., Heumos, S., Howe, K., Jain, M., Lu, T.-Y., Markello, C., Martin, F.J., Mitchell, M.W., Munson, K.M., Mwaniki, M.N., Novak, A.M., Olsen, H.E., Pesout, T., Porubsky, D., Prins, P., Sibbesen, J.A., Sirén, J., Tomlinson, C., Villani, F., Vollger, M.R., Antonacci-Fulton, L.L., Baid, G., Baker, C.A., Belyaeva, A., Billis, K., Carroll, A., Chang, P.-C., Cody, S., Cook, D.E., Cook-Deegan, R.M., Cornejo, O.E., Diekhans, M., Ebert, P., Fairley, S., Fedrigo, O., Felsenfeld, A.L., Formenti, G., Frankish, A., Gao, Y., Garrison, N.A., Giron, C.G., Green, R.E., Haggerty, L., Hoekzema, K., Hourlier, T., Ji, H.P., Kenny, E.E., Koenig, B.A., Kolesnikov, A., Korbel, J.O., Kordosky, J., Koren, S., Lee, H., Lewis, A.P., Magalhães, H., Marco-Sola, S., Marijon, P., McCartney, A., McDaniel, J., Mountcastle, J., Nattestad, M., Nurk, S., Olson, N.D., Popejoy, A.B., Puiu, D., Rautiainen, M., Regier, A.A., Rhie, A., Sacco, S., Sanders, A.D., Schneider, V.A., Schultz, B.I., Shafin, K., Smith, M.W., Sofia, H.J., Tayoun, A.N.A., Thibaud-Nissen, F., Tricomi, F.F., Wagner, J., Walenz, B., Wood, J.M.D., Zimin, A.V., Bourque, G., Chaisson, M.J.P., Flicek, P., Phillippy, A.M., Zook, J.M., Eichler, E.E., Haussler, D., Wang, T., Jarvis, E.D., Miga, K.H., Garrison, E., Marschall, T., Hall, I.M., Li, H., Paten, B.: A draft human pangenome reference. Nature 617(7960), 312–324 (2023). 10.1038/s41586-023-05896-x

[19] Guarracino, A., Buonaiuto, S., de Lima, L.G., Potapova, T., Rhie, A., Koren, S., Rubinstein, B., Fischer, C., Abel, H.J., Antonacci-Fulton, L.L., Asri, M., Baid, G., Baker, C.A., Belyaeva, A., Billis, K., Bourque, G., Carroll, A., Chaisson, M.J.P., Chang, P.-C., Chang, X.H., Cheng, H., Chu, J., Cody, S., Cook, D.E., Cook-Deegan, R.M., Cornejo, O.E., Diekhans, M., Doerr, D., Ebert, P., Ebler, J., Eichler, E.E., Eizenga, J.M., Fairley, S., Fedrigo, O., Felsenfeld, A.L., Feng, X., Flicek, P., Formenti, G., Frankish, A., Fulton, R.S., Gao, Y., Garg, S., Garrison, N.A., Giron, C.G., Green, R.E., Groza, C., Haggerty, L., Hall, I., Harvey, W.T., Haukness, M., Haussler, D., Heumos, S., Hickey, G., Hoekzema, K., Hourlier, T., Howe, K., Jain, M., Jarvis, E.D., Ji, H.P., Kenny, E.E., Koenig, B.A., Kolesnikov, A., Korbel, J.O., Kordosky, J., Lee, H., Lewis, A.P., Li, H., Liao, W.-W., Lu, S., Lu, T.-Y., Lucas, J.K., Magalhães, H., Marco-Sola, S., Marijon, P., Markello, C., Marschall, T., Martin, F.J., McCartney, A., McDaniel, J., Miga, K.H., Mitchell, M.W., Monlong, J., Mountcastle, J., Munson, K.M., Mwaniki, M.N., Nattestad, M., Novak, A.M., Nurk, S., Olsen, H.E., Olson, N.D., Paten, B., Pesout, T., Popejoy, A.B., Porubsky, D., Prins, P., Puiu, D., Rautiainen, M., Regier, A.A., Sacco, S., Sanders, A.D., Schneider, V.A., Schultz, B.I., Shafin, K., Sibbesen, J.A., Sirén, J., Smith, M.W., Sofia, H.J., Tayoun, A.N.A., Thibaud-Nissen, F., Tomlinson, C., Tricomi, F.F., Villani, F., Vollger, M.R., Wagner, J., Walenz, B., Wang, T., Wood, J.M.D., Zimin, A.V., Zook, J.M., Gerton, J.L., Phillippy, A.M., Colonna, V., Garrison, E.: Recombination between heterologous human acrocentric chromosomes. Nature 617(7960), 335–343 (2023). 10.1038/s41586-023-05976-y

[20] Crysnanto, D., Leonard, A., Pausch, H.: Comparison of methods for building pangenome graphs. In: Proceeding of 12th World Congress on Genetics Applied to Livestock Production (WCGALP) Technical and Species Orientated Innovations in Animal Breeding, and Contribution of Genetics to Solving Societal Challenges, pp. 1066–1069 (2022). Wageningen Academic Publishers

[21] Leonard, A.S., Crysnanto, D., Mapel, X.M., Bhati, M., Pausch, H.: Graph construction method impacts variation representation and analyses in a bovine super-pangenome (2022). 10.1101/2022.09.17.508368

[22] Zhou, Y., Zhang, Z., Bao, Z., Li, H., Lyu, Y., Zan, Y., Wu, Y., Cheng, L., Fang, Y., Wu, K., Zhang, J., Lyu, H., Lin, T., Gao, Q., Saha, S., Mueller, L., Fei, Z., Städler, T., Xu, S., Zhang, Z., Speed, D., Huang, S.: Graph pangenome captures missing heritability and empowers tomato breeding. Nature 606(7914), 527–534 (2022). 10.1038/s41586-022-04808-9

[23] Marçais, G., Delcher, A.L., Phillippy, A.M., Coston, R., Salzberg, S.L., Zimin, A.: MUMmer4: A fast and versatile genome alignment system. PLoS Computational Biology 14(1), 1005944 (2018). 10.1371/journal.pcbi.1005944

[24] Guarracino, A., Mwaniki, N., Marco-Sola, S., Garrison, E.: wfmash: whole-chromosome pairwise alignment using the hierarchical wavefront algorithm (2021). https://github.com/waveygang/wfmash

[25] Jain, C., Koren, S., Dilthey, A., Phillippy, A.M., Aluru, S.: A fast adaptive algorithm for computing whole-genome homology maps. Bioinformatics 34(17), 748–756 (2018). 10.1093/bioinformatics/bty597

[26] Marco-Sola, S., Moure, J.C., Moreto, M., Espinosa, A.: Fast gap-affine pairwise alignment using the wavefront algorithm. Bioinformatics (2020). 10.1093/bioinformatics/btaa777

[27] Marco-Sola, S., Eizenga, J.M., Guarracino, A., Paten, B., Garrison, E., Moreto, M.: Optimal gap-affine alignment in o(s) space. Bioinformatics 39(2) (2023). 10.1093/bioinformatics/btad074

[28] Erdős, P., Rényi, A., et al.: On the evolution of random graphs. Publ. math. inst. hung. acad. sci 5(1), 17–60 (1960)

[29] Lee, C., Grasso, C., Sharlow, M.F.: Multiple sequence alignment using partial order graphs. Bioinformatics 18(3), 452–464 (2002). 10.1093/bioinformatics/18.3.452

[30] Vaser, R., Sović, I., Nagarajan, N., Šikić, M.: Fast and accurate de novo genome assembly from long uncorrected reads. Genome Res. 27(5), 737– 746 (2017). 10.1101/gr.214270.116

[31] Gao, Y., Liu, Y., Ma, Y., Liu, B., Wang, Y., Xing, Y.: abPOA: an SIMD-based c library for fast partial order alignment using adaptive band. Bioinformatics 37(15), 2209–2211 (2020). 10.1093/bioinformatics/btaa963

[32] Vollger, M.R., Dishuck, P.C., Harvey, W.T., DeWitt, W.S., Guitart, X., Goldberg, M.E., Rozanski, A.N., Lucas, J., Asri, M., Abel, H.J., Antonacci-Fulton, L.L., Baid, G., Baker, C.A., Belyaeva, A., Billis, K., Bourque, G., Buonaiuto, S., Carroll, A., Chaisson, M.J.P., Chang, P.-C., Chang, X.H., Cheng, H., Chu, J., Cody, S., Colonna, V., Cook, D.E., Cook-Deegan, R.M., Cornejo, O.E., Diekhans, M., Doerr, D., Ebert, P., Ebler, J., Eizenga, J.M., Fairley, S., Fedrigo, O., Felsenfeld, A.L., Feng, X., Fischer, C., Flicek, P., Formenti, G., Frankish, A., Fulton, R.S., Gao, Y., Garg, S., Garrison, E., Garrison, N.A., Giron, C.G., Green, R.E., Groza, C., Guarracino, A., Haggerty, L., Hall, I.M., Haukness, M., Haussler, D., Heumos, S., Hickey, G., Hourlier, T., Howe, K., Jain, M., Jarvis, E.D., Ji, H.P., Kenny, E.E., Koenig, B.A., Kolesnikov, A., Korbel, J.O., Kordosky, J., Koren, S., Lee, H., Li, H., Liao, W.-W., Lu, S., Lu, T.-Y., Lucas, J.K., Magalhães, H., Marco-Sola, S., Marijon, P., Markello, C., Marschall, T., Martin, F.J., McCartney, A., McDaniel, J., Miga, K.H., Mitchell, M.W., Monlong, J., Mountcastle, J., Mwaniki, M.N., Nattestad, M., Novak, A.M., Nurk, S., Olsen, H.E., Olson, N.D., Paten, B., Pesout, T., Phillippy, A.M., Popejoy, A.B., Prins, P., Puiu, D., Rautiainen, M., Regier, A.A., Rhie, A., Sacco, S., Sanders, A.D., Schneider, V.A., Schultz, B.I., Shafin, K., Sibbesen, J.A., Sirén, J., Smith, M.W., Sofia, H.J., Tayoun, A.N.A., Thibaud-Nissen, F., Tomlinson, C., Tricomi, F.F., Villani, F., Vollger, M.R., Wagner, J., Walenz, B., Wang, T., Wood, J.M.D., Zimin, A.V., Zook, J.M., Munson, K.M., Lewis, A.P., Hoekzema, K., Logsdon, G.A., Porubsky, D., Paten, B., Harris, K., Hsieh, P., Eichler, E.E.: Increased mutation and gene conversion within human segmental duplications. Nature 617(7960), 325–334 (2023). 10.1038/s41586-023-05895-y

[33] Heumos, S., Guarracino, A., Schmelzle, J.-N.M., Li, J., Zhang, Z., Hagmann, J., Nahnsen, S., Prins, P., Garrison, E.: Pangenome graph layout by path-guided stochastic gradient descent. bioRxiv (2023) https://www.biorxiv.org/content/early/2023/10/17/2023.09.22.558964.full.pdf. 10.1101/2023.09.22.558964

[34] Ewels, P., Magnusson, M., Lundin, S., Käller, M.: Multiqc: summarize analysis results for multiple tools and samples in a single report. Bioinformatics 32(19), 3047 (2016). 10.1093/bioinformatics/btw354

[35] Paten, B., Eizenga, J.M., Rosen, Y.M., Novak, A.M., Garrison, E., Hickey, G.: Superbubbles, ultrabubbles, and cacti. Journal of Computational Biology 25(7), 649–663 (2018). 10.1089/cmb.2017.0251

[36] Garrison, E., Kronenberg, Z.N., Dawson, E.T., Pedersen, B.S., Prins, P.: A spectrum of free software tools for processing the VCF variant call format: vcflib, bio-vcf, cyvcf2, hts-nim and slivar. PLoS Computational Biology 18(5), 1009123 (2022). 10.1371/journal.pcbi.1009123

[37] Heumos, S., Hanssen, F., Heumos, L., Guarracino, A., Heuer, M., Ehmele, P., Prins, P., Garrison, E., Nahnsen, S.: nf-core/pangenome (2024). 10.5281/zenodo.8202637

[38] Clark, K., Karsch-Mizrachi, I., Lipman, D.J., Ostell, J., Sayers, E.W.: Genbank. Nucleic Acids Research 44(D1), 67–72 (2015). 10.1093/nar/gkv1276

[39] O’Donnell, S., Yue, J.-X., Saada, O.A., Agier, N., Caradec, C., Cokelaer, T., De Chiara, M., Delmas, S., Dutreux, F., Fournier, T., Friedrich, A., Kornobis, E., Li, J., Miao, Z., Tattini, L., Schacherer, J., Liti, G., Fischer, G.: Telomere-to-telomere assemblies of 142 strains characterize the genome structural landscape in saccharomyces cerevisiae. Nature Genetics 55(8), 1390–1399 (2023). 10.1038/s41588-023-01459-y

[40] Liu, Y., Du, H., Li, P., Shen, Y., Peng, H., Liu, S., Zhou, G.-A., Zhang, H., Liu, Z., Shi, M., Huang, X., Li, Y., Zhang, M., Wang, Z., Zhu, B., Han, B., Liang, C., Tian, Z.: Pan-genome of wild and cultivated soybeans. Cell 182(1), 162–17613 (2020). 10.1016/j.cell.2020.05.023

[41] Chu, J.S.-C., Peng, B., Tang, K., Yi, X., Zhou, H., Wang, H., Li, G., Leng, J., Chen, N., Feng, X.: Eight soybean reference genome resources from varying latitudes and agronomic traits. Scientific Data 8(1) (2021). 10.1038/s41597-021-00947-2

[42] Zhou, Y., Zhang, Z., Bao, Z., Li, H., Lyu, Y., Zan, Y., Wu, Y., Cheng, L., Fang, Y., Wu, K., Zhang, J., Lyu, H., Lin, T., Gao, Q., Saha, S., Mueller, L., Fei, Z., Städler, T., Xu, S., Zhang, Z., Speed, D., Huang, S.: Graph pangenome captures missing heritability and empowers tomato breeding. Nature 606(7914), 527–534 (2022). 10.1038/s41586-022-04808-9

[43] Llamas, B., Narzisi, G., Schneider, V., Audano, P.A., Biederstedt, E., Blauvelt, L., Bradbury, P., Chang, X., Chin, C.-S., Fungtammasan, A., Clarke, W.E., Cleary, A., Ebler, J., Eizenga, J., Sibbesen, J.A., Markello, C.J., Garrison, E., Garg, S., Hickey, G., Lazo, G.R., Lin, M.F., Mahmoud, M., Marschall, T., Minkin, I., Monlong, J., Musunuri, R.L., Sagayaradj, S., Novak, A.M., Rautiainen, M., Regier, A., Sedlazeck, F.J., Siren, J., Souilmi, Y., Wagner, J., Wrightsman, T., Yokoyama, T.T., Zeng, Q., Zook, J.M., Paten, B., Busby, B.: A strategy for building and using a human reference pangenome. F1000Res 8, 1751 (2021). 10.12688/f1000research.19630.2

[44] Traag, V.A., Waltman, L., van Eck, N.J.: From louvain to leiden: guaranteeing well-connected communities. Scientific Reports 9(1) (2019). 10.1038/s41598-019-41695-z

[45] Kosugi, S., Momozawa, Y., Liu, X., Terao, C., Kubo, M., Kamatani, Y.: Comprehensive evaluation of structural variation detection algorithms for whole genome sequencing. Genome Biology 20(1) (2019). 10.1186/s13059-019-1720-5

[46] Poplin, R., Chang, P.-C., Alexander, D., Schwartz, S., Colthurst, T., Ku, A., Newburger, D., Dijamco, J., Nguyen, N., Afshar, P.T., Gross, S.S., Dorfman, L., McLean, C.Y., DePristo, M.A.: A universal snp and small-indel variant caller using deep neural networks. Nature Biotechnology 36(10), 983–987 (2018). 10.1038/nbt.4235

[47] Ondov, B.D., Treangen, T.J., Melsted, P., Mallonee, A.B., Bergman, N.H., Koren, S., Phillippy, A.M.: Mash: fast genome and metagenome distance estimation using minhash. Genome Biology 17(1) (2016). 10.1186/s13059-016-0997-x

[48] Paten, B., Earl, D., Nguyen, N., Diekhans, M., Zerbino, D., Haussler, D.: Cactus: Algorithms for genome multiple sequence alignment. Genome Research 21(9), 1512–1528 (2011). 10.1101/gr.123356.111

[49] Doerr, D.: Panacus: A Counting Tool for Pangenome Graph (2023). https://github.com/marschall-lab/panacus

[50] Rasko, D.A., Rosovitz, M.J., Myers, G.S.A., Mongodin, E.F., Fricke, W.F., Gajer, P., Crabtree, J., Sebaihia, M., Thomson, N.R., Chaudhuri, R., Henderson, I.R., Sperandio, V., Ravel, J.: The pangenome structure ofescherichia coli: Comparative genomic analysis ofe. colicommensal and pathogenic isolates. Journal of Bacteriology 190(20), 6881–6893 (2008). 10.1128/jb.00619-08

[51] Lin, T., Zhu, G., Zhang, J., Xu, X., Yu, Q., Zheng, Z., Zhang, Z., Lun, Y., Li, S., Wang, X., Huang, Z., Li, J., Zhang, C., Wang, T., Zhang, Y., Wang, A., Zhang, Y., Lin, K., Li, C., Xiong, G., Xue, Y., Mazzucato, A., Causse, M., Fei, Z., Giovannoni, J.J., Chetelat, R.T., Zamir, D., Städler, T., Li, J., Ye, Z., Du, Y., Huang, S.: Genomic analyses provide insights into the history of tomato breeding. Nature Genetics 46(11), 1220–1226 (2014). 10.1038/ng.3117

[52] Ou, S., Su, W., Liao, Y., Chougule, K., Agda, J.R.A., Hellinga, A.J., Lugo, C.S.B., Elliott, T.A., Ware, D., Peterson, T., Jiang, N., Hirsch, C.N., Hufford, M.B.: Benchmarking transposable element annotation methods for creation of a streamlined, comprehensive pipeline. Genome Biology 20(1) (2019). 10.1186/s13059-019-1905-y

[53] Robinson, J.T., Thorvaldsdóttir, H., Winckler, W., Guttman, M., Lander, E.S., Getz, G., Mesirov, J.P.: Integrative genomics viewer. Nature Biotechnology 29(1), 24–26 (2011). 10.1038/nbt.1754

